# Dynamic BH3 profiling predicts clinical outcomes in acute myeloid leukemia

**DOI:** 10.1101/2025.10.14.682414

**Authors:** Emma-Jayne Minihane, Leah Krotee, Jason Wu, Lindsey Thornton, Geoffrey Fell, Sonja Herter, Jeremy Ryan, Amulya Jakkani, Kate Marinchev, Andrew A. Lane, Marlise R. Luskin, Eric S. Winer, Evan C. Chen, Shai Shimony, Martha Wadleigh, Rahul Vedula, Virginia O. Volpe, Christopher Reilly, Ilene Galinsky, Daniel J. DeAngelo, Richard M. Stone, Donna S. Neuberg, Anthony Letai, Jacqueline S. Garcia

**Affiliations:** Department of Medical Oncology, Dana-Farber Cancer Institute, Boston, MA; Department of Data Science, Dana-Farber Cancer Institute, Boston, MA

**Keywords:** Drug-induced apoptotic priming

## Abstract

Predictive biomarkers can potentially meet the need for improved drug assignment in acute myeloid leukemia (AML). Fewer than half of AML patients have actionable mutations: consequently, targeted therapy achieves remission in only a fraction of those who have them. Dynamic BH3 Profiling (DBP), a functional assay, can measure changes in *ex vivo* drug-induced apoptotic priming in multiple cancers. To assess the feasibility and predictive capacity of DBP in AML, we prospectively tested DBP using a fixed-drug panel in myeloblasts from 92 patients. We generated a database combining genetic and functional annotation. Established AML clinical and genetic prognostic characteristics were associated with drug-induced apoptotic priming. We observed distinct *inter*patient sensitivities to single drugs or combinations with the BCL2-inhibitor venetoclax, and *intra*patient apoptotic priming differences based on CD123-expression within distinct cell subpopulations. DBP further predicted the likelihood of remission to chemotherapy and targeted agents, supporting its use to identify optimal personalized therapy.

**Statement of significance:** Dynamic BH3 profiling provides patient-specific drug vulnerability data in real-time to inform prognosis and therapy selection.

**Key takeaways:**

1. Dynamic BH3 profiling can be performed on bone marrow and leukemic blood from AML patients in 48 hours.
2. Known clinical prognostic factors associate with drug-induced apoptotic priming in AML.
3. Drug-induced apoptotic priming identifies drug vulnerabilities in individual patients and predicts clinical response to chemotherapy and small molecule inhibitors.

## Introduction

Acute myeloid leukemia (AML) is a heterogenous hematologic cancer that is lethal if untreated. Despite major advances in the therapeutic landscape for AML over the last decade, most patients experience disease relapse and death. Treatment failure remains the primary cause of death. Less than 20% of patients who are refractory to conventional cytotoxic induction chemotherapy or relapse after allogeneic transplant are alive at 1-year [1]. The development of targeted therapies for AML with *IDH1/2, FLT3-*ITD/TKD, and *KMT2A* rearrangement and less intensive chemotherapy regimens have increased treatment options for patients experiencing treatment failure, and for patients ≥ 75 years or younger patients with significant co-morbidities. Despite an expanding therapeutic portfolio of targeted therapies, most patients do not have an actionable target or do not respond durably to genetically directed single-agent targeted therapy (∼30% rate of full complete remission) in the relapsed or refractory (R/R) setting across therapies, and long-term remissions require successful allogeneic transplant. Thus, while genetic assessment has become the focus of AML classification at diagnosis and at relapse, there is an urgent clinical need for additional biomarkers capable of informing therapeutic selection [2–4].

This need prompted our investigation of predictive biomarkers to formulate a depiction of drug vulnerabilities in order to optimize future clinical trial design. While a growing number of *ex vivo* studies in AML have been exploring the utility of functional assays in the preclinical settings or have been applied retrospectively [5–8], functional assays as predictive biomarkers are not yet integrated into routine cancer care. With an expansive treatment arsenal, we set out to develop a predictive biomarker that would steer individual patients towards effective therapies and away from those that are futile.

We previously tested baseline (routine) BH3 profiling, which is a functional assay, on myeloblasts collected from pre-treatment AML patients. Routine BH3 profiling measures cytochrome *c* retention following mitochondrial exposure to synthetic BH3 peptides [9–11]. The lower cytochrome *c* is retained (indicating cytochrome *c* release), the greater the apoptotic priming. We found that apoptotic priming was associated with sensitivity to conventional induction chemotherapy [9]. In dynamic BH3 profiling (DBP), BH3 profiling is preceded by treatment of myeloblasts to drug(s) for a short incubation period (16-24 hours) so that drug-induced apoptotic priming can be measured and compared to a drug-untreated control (DMSO) in viable cells (termed *delta priming*). An advantage of DBP over baseline BH3 profiling is that it can detect patient-specific drug vulnerabilities across a wider spectrum of agents. In AML PDX models, DBP identified drug sensitivities even in the context of AML resistant/refractory to the BH3-mimetic venetoclax [12]. In retrospective studies, DBP performed on viably frozen AML marrow samples predicted achievement of complete remission (CR) to azacitidine/venetoclax (aza/ven) combinations or gilteritinib monotherapy [12, 13]. DBP predicted complete remission achievement in two phase 1 clinical trials evaluating lenalidomide followed by salvage chemotherapy in R/R AML [14]. We further performed routine BH3 profiling of paired leukemic blood samples collected from patients before and after single-agent lenalidomide lead-in treatment prior to chemotherapy. This demonstrated *in vivo* lenalidomide-induced apoptotic priming, providing proof-of-concept that lenalidomide sensitized myeloblasts to subsequent chemotherapy. Since DBP detects apoptotic signaling induced by any treatment, it has the potential to broadly guide real-time therapeutic decisions. Our assay is of particular value as it measures a specific apoptotic signal within single cells and can identify priming differences within cell subpopulations as well as bulk populations all within a minimal cell culture time (∼16 hours).

AML exhibits great heterogeneity within (*intra*patient) and between (*inter*patient) patients. While genetic heterogeneity influences patient outcomes [3, 15, 16] less is understood about functional heterogeneity especially as it relates to predicting treatment response. Choice of a therapy beyond a mutationally-guided algorithm would be advantageous. In the present study, we prospectively performed 101 DBPs using fresh bone marrow aspirate from 92 unique AML patients with a pre-specified panel of drugs. We then integrated functional data from this *ex vivo* drug sensitivity assay with molecular and clinical factors, establishing an ‘AML DBP database’. From this large comprehensive dataset, we tested its clinical utility against validated AML prognostic factors and its predictive power for patients who later went on to receive chemotherapy that was included in the fixed drug-panel. These findings suggest that the addition of DBP to our current standard diagnostic work-up has potential for prospective use to inform optimal treatment selection in AML.

## Results

### Heterogeneous drug-induced apoptotic priming is observed in patient AML myeloblasts

To investigate biological heterogeneity among patients with AML, we established a bedside-to-bench research pipeline to collect bone marrow aspirate samples from AML patients at time of routine disease assessment. Bone marrow aspirates were collected at the time of new diagnosis or relapse to undergo DBP (**Fig. 1a**). We performed DBP in 101 freshly collected patient samples (59 aspirate, 5 leukapheresis, and 37 leukemic blood) from 92 unique patients representing a broad spectrum of AML subtypes in untreated (N=58), and R/R (N=34) disease states, but none were paired diagnostic/relapse samples (**Fig. 1b**, **Table 1, Supplemental Table 1**). Biological duplicates (N=9) included 3/9 at unique R/R timepoints, 4/9 collected from different sources (matched blood/marrow), and 2/9 same timepoint/source but independent collection and separate DBP run. DBP data were not made available to clinicians as data optimization was on-going. We subsequently created an AML DBP database, including detailed clinical, pathological and molecular information along with DBP results. Treatment in clinic was administered to 79/92 (86%) patients. Clinical response assessments were recorded according to European LeukemiaNet (ELN) 2022 AML response criteria in 70/79 (89%) evaluable patients who had follow-up and subsequent disease assessment including bone marrow biopsy.

**Figure 1.**
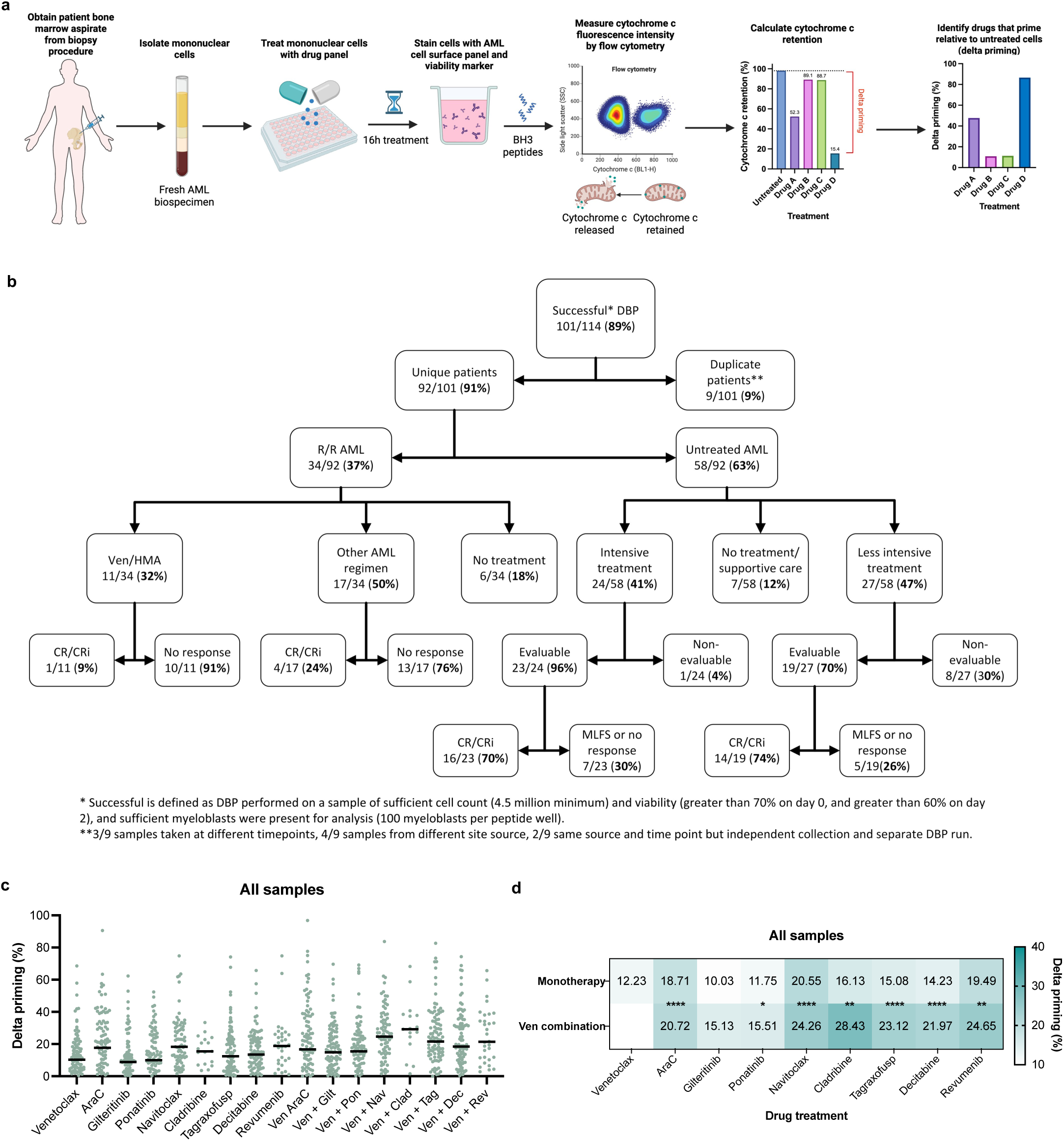
Evidence for heterogeneous drug-induced apoptotic priming among AML patients. (**a**) Overview of the dynamic BH3 profiling assay from sample collection to data assessment. Turnaround time is 48-hours from sample arrival to data analysis. (**b**) Consort diagram to demonstrate feasibility of DBP in AML and clinical responses to non-DBP assigned therapy after sample collection. Additional information can be found in **Supplemental Table 1**. (**c**) Mean delta priming values calculated from results of 101 samples from 92 unique patients. 10 nM BIM peptide was used. Each dot represents the average of 4-6 peptide replicates for a single patient sample. Black line represents median. (**d**) Mean delta priming for each monotherapy is compared to mean delta priming for each venetoclax-combination. Significance determined by one-way ANOVA and Tukey’s multiple comparisons test.

**Table 1.**
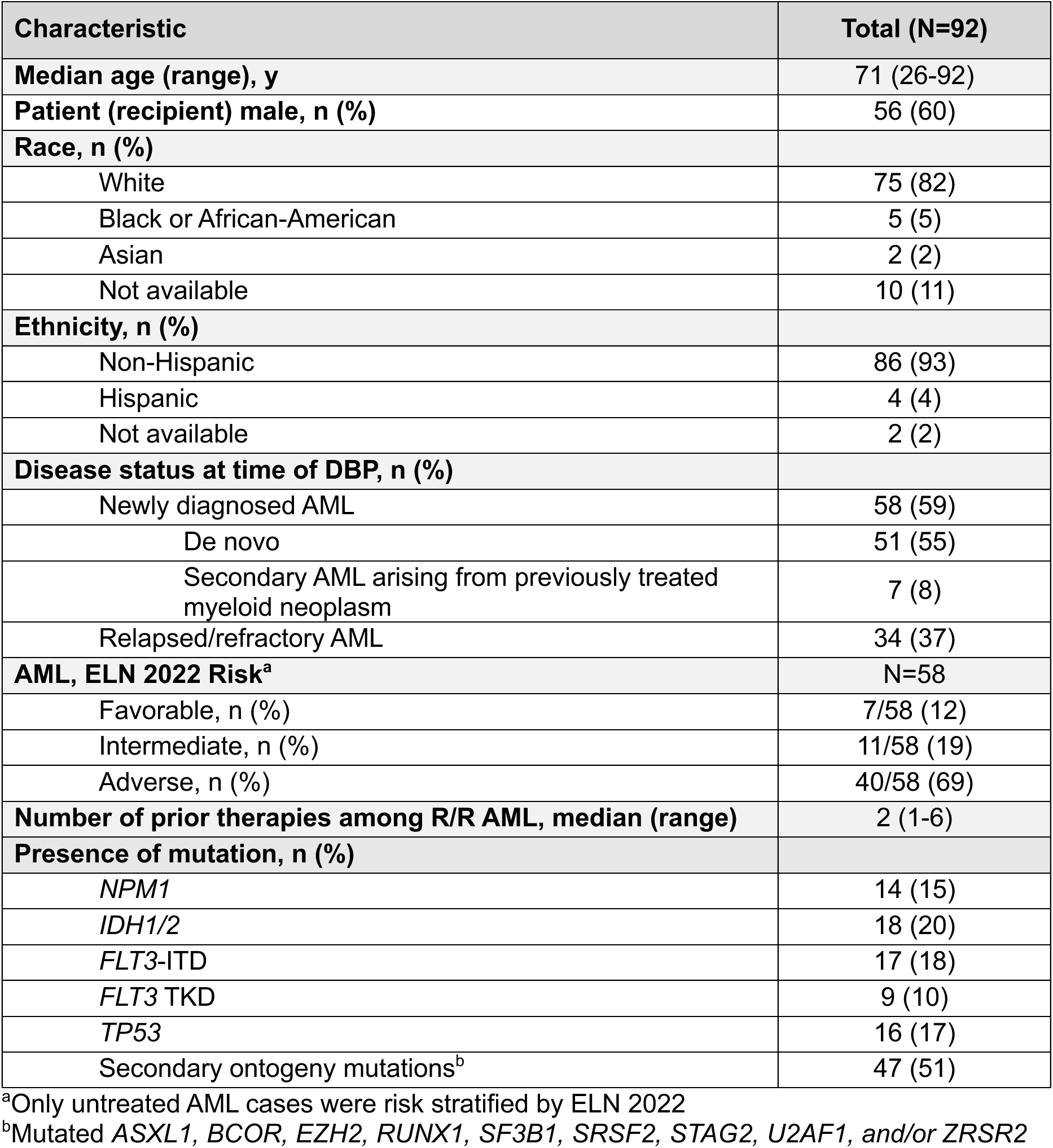
Patient- and disease-related characteristics.

Feasibility (or ‘successful DBP’) was defined as meeting three parameters: sufficient cell count (∼4.5 million to test proposed drug conditions with 25,000 cells per peptide well), cell viability (70% on day 0 and >60% on day 2), and at least 100 viable myeloblasts per peptide well upon analysis. Feasibility of the assay was determined to be 88.8%, with assay failures mostly occurring due to equipment errors. Over the 34 months of data collection, 13 technical errors occurred (13/114) with 11/13 occurring in the first 6 months. The median time from biospecimen receipt to data output was 2 days (range, 2-5). Only 15/101 samples required 4-5 days to read out, largely due to holiday weekend or machine downtime. As a conceivable application of DBP includes consideration of practical AML therapies, we designed a drug-panel consisting of a broad range of agents with available safety and biological activity either alone or in combination with venetoclax in acute leukemia patients [17–25]; thus potentially clinically actionable. As proof-of-concept, we designed a fixed drug-panel by focusing on agents with distinct mechanisms of action, including each of the following as monotherapy venetoclax, cladribine, cytarabine, decitabine, gilteritinib, navitoclax, ponatinib, revumenib, or tagraxofusp, and each in combination with venetoclax. Further, we anticipated some patients might have received some of these proposed therapies on the drug-panel in the clinic, which would enable retrospective analysis of DBP-predicted responses.

A useful biomarker would be expected to show *inter*patient heterogeneity, and we observed differential drug-induced priming responses by DBP across our cohort (**Fig. 1c**). In 7/8 of the drug-combinations, venetoclax increased overall drug-induced apoptotic priming compared to that observed with monotherapy regardless of disease status (relapsed or newly diagnosed), with the exception being the FLT3-inhibitor gilteritinib (**Fig. 1d; Fig. S1a**). DBP further identified myeloblasts that did not show increased drug-induced priming with the addition of venetoclax based on minimal to no increase in drug-induced apoptotic priming, which suggests low likelihood of drug sensitivity.

Another major advantage of DBP is the rapidity of signal from early measurement of apoptotic induction via cytochrome *c* release at the level of mitochondrial outer membrane permeabilization (MOMP) in response to BH3 peptides, which is detectable at 16-hours of drug treatment (**Fig. S1b**). However, the Annexin V assay, a measure of completed cell death, provided a useful discriminating signal only after a 72-hour incubation (**Fig. S1c-d**). The relatively brief incubation period required for DBP thus minimizes the artifacts inherent in *ex vivo* culture time which may cause selection, adaptation, or spontaneous cell death.

### Drug-induced apoptotic priming associates with established AML prognostic factors

Factors predictive of prognosis in AML and response to intensive therapy include age and the European LeukemiaNet (ELN) 2022 risk classification, which is based on the presence or absence of certain cytogenetics and genetic mutations [3]. We hypothesized that if DBP was returning clinically meaningful data, higher delta priming would correlate with good prognostic factors in newly diagnosed AML, and lower delta priming with poor prognostic factors. To address this question, we pooled delta priming data for all tested drugs, providing 1340 individual *ex vivo* drug-test results from 58 unique untreated AML patients. We observed that myeloblasts from AML patients ≥60 years had overall reduced drug-induced apoptotic priming compared to those from patients <60 years (**Fig. 2a**; P<0.0001). Significantly reduced apoptotic priming for patients with ELN adverse risk was observed when compared to patients with Favorable (P<0.0001) or Intermediate risk (P<0.0001) (**Fig. 2b**). To determine if the magnitude of apoptotic priming in each risk group correlated with improved progression-free survival (PFS), we separated DBP test results by above or below the median delta priming and generated Kaplan-Meier curves after a median follow-up of 181 days (range, 3-910 days) (**Fig. 2c-e**). Patients with increased delta priming above the pooled median trended towards prolonged PFS for those with Favorable ELN risk (**Fig. 2c**; P=0.02) but did not significantly improve PFS for those with Intermediate ELN risk where there is the most disease heterogeneity (**Fig. 2d**; P=0.15). In the Adverse risk group, no difference was observed (**Fig. 2e**; P=0.23).

**Figure 2.**
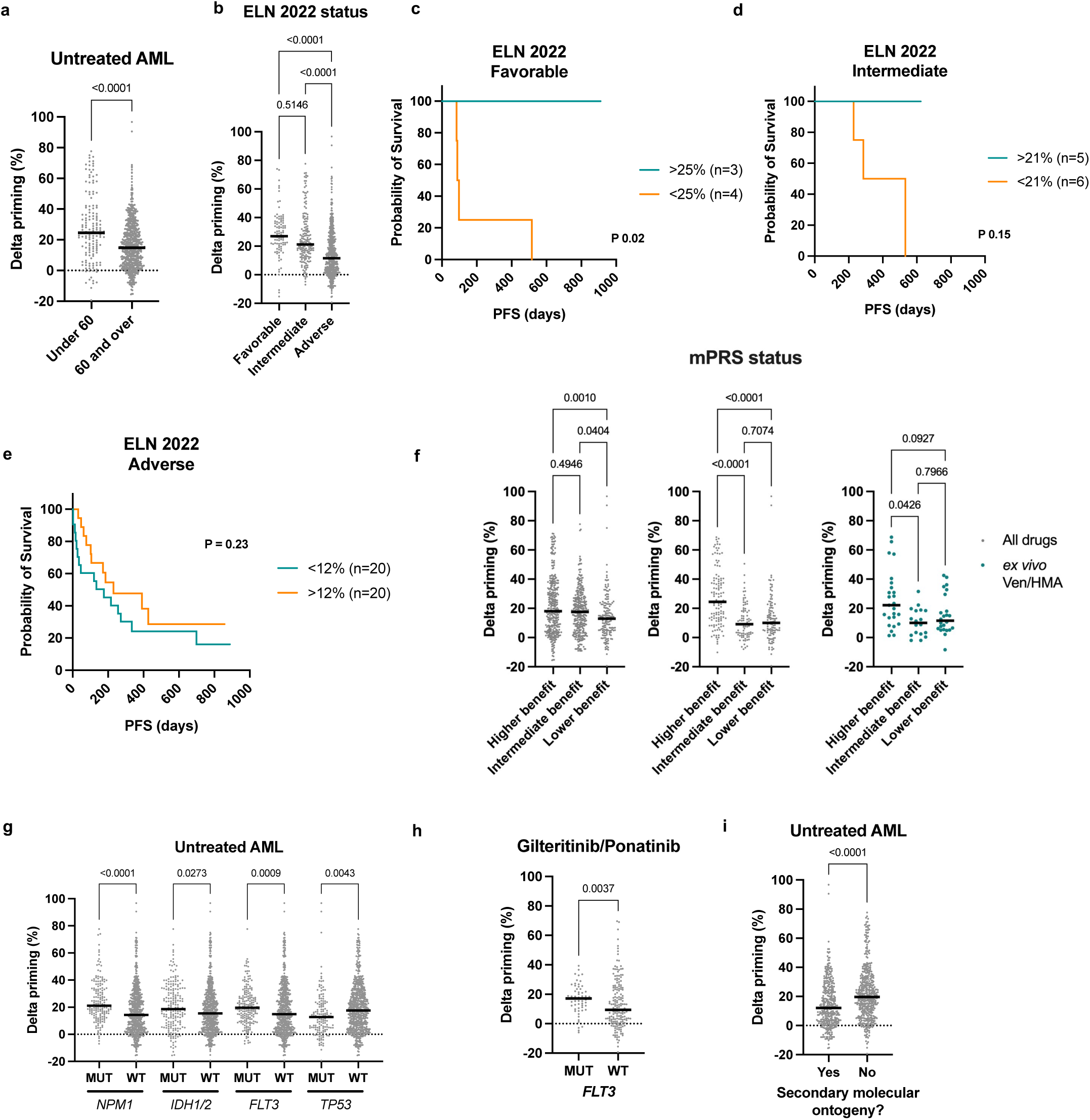
Drug-induced priming patterns associate with known prognostic features in AML. Among 58 untreated AML cases: (**a**) Comparison of pooled delta priming data in patients under the age of 60 years (n=152) or ≥60 years (n=675; one-tailed Mann-Whitney test). (**b**) Pooled delta priming data in AML patients stratified into ELN 2022 risk groups (Favorable n=98, Intermediate n=168, Adverse n=518; one-way ANOVA with Tukey’s multiple comparisons). (**c-e**) Samples were divided into above or below median delta priming for each ELN 2022 risk group and a Kaplan-Meier plot of PFS was generated for each ELN classification including (**c**) Favorable risk patients (N=7; median 25%), (**d**) Intermediate risk patients (N=11; median 21%), and (**e**) Adverse risk patients (N=40; median 12%) (one-tailed log-rank test). (**f**) (left) Delta priming data for untreated AML samples grouped by mPRS status (Higher benefit n=336, Intermediate benefit n=334, Lower benefit n=157). (middle) Pooled delta priming data from 20 untreated AML patients who went on to receive HMA-based therapy in the clinic after DBP collection were grouped by mPRS status (Higher benefit n=120, Intermediate benefit n=82, Lower benefit n=109; (right) Delta priming data for the same 20 untreated AML patient grouped by mPRS status based on *ex vivo* treatment with Ven or Decitabine or Ven/Dec combination (Kruskal-Wallis test). (**g**) Pooled delta priming data from untreated AML patients comparing priming in myeloblasts with mutated (Mut) versus wildtype (WT) *NPM1* (Mut n=167, WT n=660), *IDH1/2* (Mut n=196, WT n=631), *FLT3-*ITD/TKD (Mut n=194, WT n=620) and *TP53* (Mut = 144, WT = 683) (Kruskal-Wallis with Dunn’s multiple comparisons). (**h**) Comparison of pooled delta priming for *FLT3* MUT (n=56) and *FLT3* WT (n=179) AML samples treated *ex vivo* with gilteritinib or ponatinib (one-tailed Mann-Whitney test). (**i**) Comparison of delta priming in AML patients with (n=366) or without (n=461) secondary molecular ontogeny mutations (one-tailed Mann-Whitney test).

With increased use of less-intensive therapies in AML, the molecular prognostic risk signature (mPRS) using four genes was proposed to stratify outcomes for patients who receive HMA/ven, where absence of mutations in the four genes indicates *higher benefit*, presence of mutation in *TP53* indicates *lower benefit*, and presence of mutation in *FLT3*-ITD, *KRAS*, or *NRAS* indicates *intermediate benefit* [26]. We hypothesized that higher apoptotic priming would be observed among the *higher benefit* patients. Among the untreated AML samples, we observed higher delta priming in myeloblasts from the *higher benefit* and *intermediate benefit* groups compared to those from the *lower benefit* samples (**Fig. 2f, left,** P<0.04). We further asked if this would also apply to patients that received HMA/ven-based therapy in the clinic, aligning with clinical outcomes. In our database, we identified 20 patients with untreated AML who received HMA/ven-based therapy after DBP sample collection and stratified them by mPRS status. As hypothesized, the median delta priming was significantly higher in the *higher benefit* when compared to the *lower* and *intermediate* benefit groups (**Fig. 2f, middle;** P<0.043). To determine the extent mPRS and DBP prediction was applicable beyond HMA/ven, we pooled all drug treatments together in the same set of patients (N=20; **Fig. 2f, right,** P<0.0001). We observed that mPRS also associates with apoptotic priming in response to a broad panel of drugs.

To further investigate the correlation of specific genetic mutations on drug-induced priming, we focused on targetable mutations or those associated with chemosensitivity. The presence of actionable mutations in AML provides the opportunity for the use of targeted therapies, e.g. gilteritinib in *FLT3*-mutated AML, but clinical outcomes with these targeted therapies suggest additional biomarkers are needed to predict clinical response. Mutations in *NPM1* confer chemosensitivity and favorable outcomes, especially without a *FLT3*-ITD co-mutation [27, 28]. Compared to AML cases wild-type for the mutation, we observed significantly increased drug-induced apoptotic priming among pooled DBP in myeloblasts with mutations in *NPM1* (P<0.0001), *IDH1*/2 (P=0.0273), and *FLT3* (P=0.0009) (**Fig. 2g**). In contrast, as predicted, significantly decreased drug-induced apoptotic priming was observed in myeloblasts from *TP53*-mutated pooled samples compared to wild-type (P=0.0043). Despite overall increased delta priming among the *NPM1, IDH1/2* and *FLT3-*mutated pooled samples, we saw minimal differences in mean delta priming with most individual drugs within mutated subgroups (**Fig. S2a**). However, tyrosine kinase inhibitors, such as gilteritinib (*FLT3*-selective) and ponatinib (multi-targeted), yielded increased delta priming in *FLT3*-mutated samples versus wildtype cases (**Fig. 2h**; P=0.004). A significant minority of *FLT*3 wild-type samples exhibited an apoptotic priming response to these TKIs, congruent with the clinical observation that patients with wild-type *FLT3* have a detectable level of clinical response to these drugs, though an available clinical biomarker to identify them has been lacking [29, 30].

We next asked if molecular ontogeny influenced drug-induced apoptotic priming. Myelodysplasia-related defining mutations or secondary ontogeny include those in *ASXL1, BCOR, EZH2, RUNX1, SF3B1, SRSF2, STAG2, U2AF1, ZRSR2*, and are associated with poorer prognosis compared to *de novo* AML leading to its recent adverse risk assignment in ELN 2022 [3, 15]. In agreement with the clinical data, we observed overall reduced drug-induced apoptotic priming in the presence versus absence of mutations associated with secondary molecular ontogeny (**Fig. 2i**; P<0.0001).

### DBP identifies *ex vivo* drug sensitivities despite relapsed/refractory AML status

Upon disease relapse, reduced overall apoptotic priming is a resistance mechanism across anti-cancer therapies [9, 13]. Based on our prior data [12], we hypothesized that the overall heterogeneity of apoptotic priming in response to our drug panel would be reduced among R/R AML samples compared to that observed in previously untreated AML samples. Using pooled delta priming responses, delta priming was overall reduced in R/R versus untreated AML (**Fig. 3a left,** P=0.002), however the overall variation of delta priming was slightly higher in the R/R AML pooled dataset (**Fig. 3a right;** ratio 0.8:1). Drug-induced apoptotic priming to *ex vivo* combination venetoclax and cytarabine (AraC) treatment patients with R/R AML was reduced compared untreated AML; this combination has less clinical activity and durability in the advanced disease setting (**Fig. 3b**; P=0.018). Next, we asked if there were differences in drug-induced priming among R/R AML patients based on prior venetoclax exposure. We found an overall increased delta priming despite prior venetoclax exposure versus none (**Fig. 3c**; P=0.0076). No differences in delta priming were observed with monotherapies based on prior venetoclax exposure status (P=0.810), suggesting that venetoclax can still promote apoptosis with new partner agents despite prior use.

**Figure 3.**
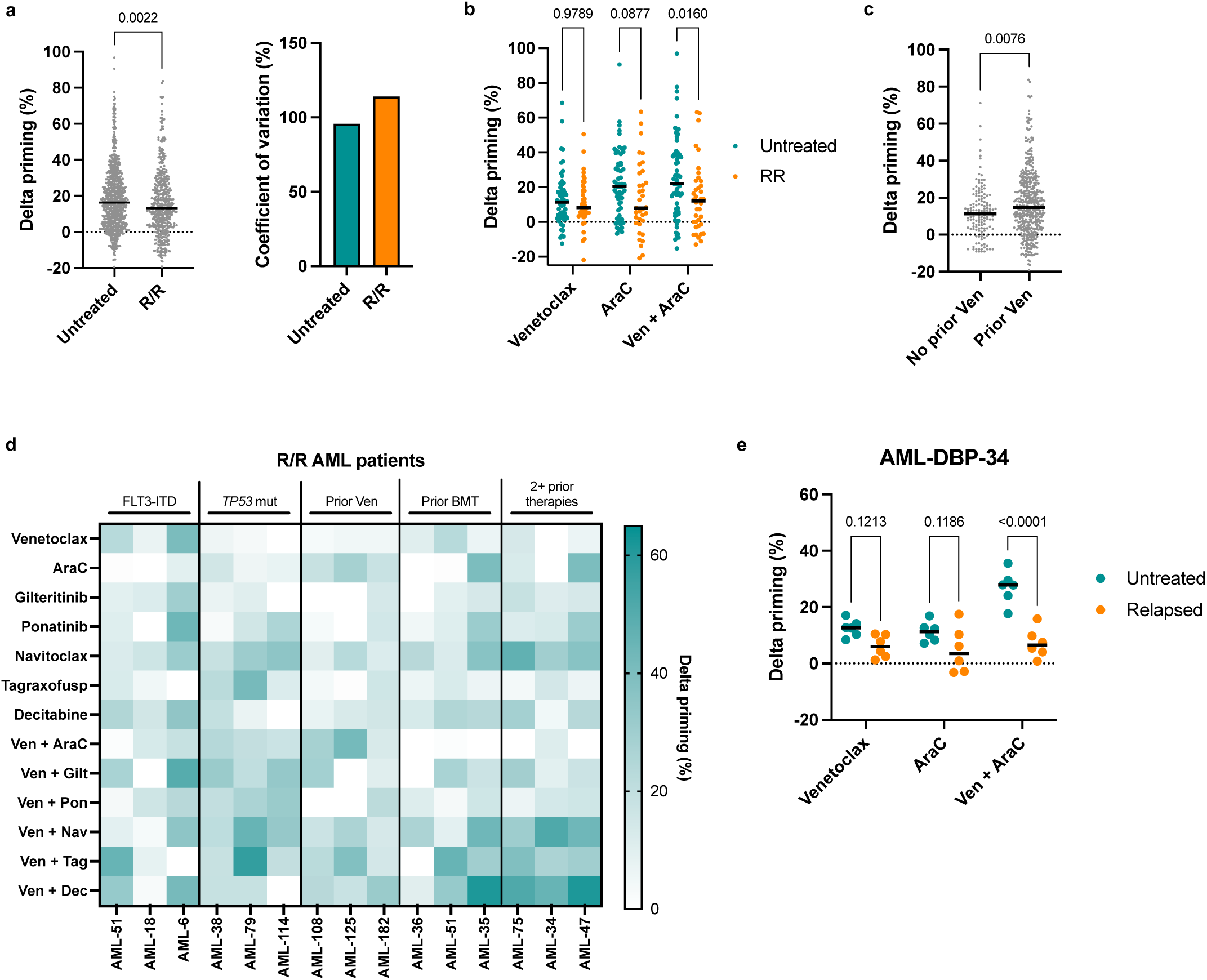
DBP identifies drug-induced apoptotic priming changes even in the relapsed/refractory setting. (**a**) (left) Comparison of pooled delta priming data from myeloblasts from untreated (n=827) and R/R, n=423 patients (two-tailed Mann Whitney test), (right) coefficient of variation for each group is compared. (**b**) Delta priming in response to venetoclax, AraC, or the combination was compared between untreated and R/R samples (two-way ANOVA with Tukey’s multiple comparisons). (**c**) Comparison of pooled delta priming for R/R samples with no prior venetoclax (ven) treatment history versus prior venetoclax treatment (two-tailed Mann Whitney test). (**d**) Mean delta priming for individual patients representing common AML subgroups is compared. Data are mean of 6 BIM 10nM peptide replicates. (**e**) Delta priming in response to venetoclax, AraC, and combination venetoclax-AraC in myeloblasts from paired untreated and relapsed samples from AML patient DBP-34 (two-way ANOVA with Tukey’s multiple comparisons).

Across common clinical subgroups among R/R AML patients, we observed broad drug-induced priming (**Fig. 3d**). We hypothesized that drug-induced priming would be reduced at the time of relapse compared to diagnosis. Sufficient cells to test three drug conditions from paired bone marrow samples collected at diagnosis (viably frozen cells from the tissue bank) and relapse (collected fresh) (47 months apart) were available for one patient. We observed no differences in baseline priming in myeloblasts at diagnosis or relapse using BIM peptide alone with DMSO/untreated (90% cytochrome c retention). Compared to myeloblasts obtained at diagnosis, we observed a significant delta priming decrease in those at relapse with combination venetoclax and AraC, and a numerical decrease with venetoclax or AraC as monotherapy, which would have suggested a low likelihood of response (**Fig. 3e**, P<0.0001). These findings are consistent with the patient’s clinical course: initial AraC-based therapy induced remission, but upon relapse did not respond to subsequent AraC/clofarabine nor after that aza/ven-based therapy.

### DBP captures antigen-positive drug-induced priming within subpopulations in AML

The level of the stem cell antigen CD123 expression on myeloblasts correlates with poorer AML outcomes [31]. The CD123-directed cytotoxin tagraxofusp is approved for use in blastic plasmacytoid dendritic cell neoplasm, and preliminary results show promising safety and activity in combination with aza/ven as a triplet regimen in AML[23, 32]. As our DBP assay AML antibody panel includes CD123, we asked if drug-induced priming by tagraxofusp also correlated with CD123-expressing myeloblasts. In our DBP database, 9/101 had unknown CD123 status (not available on clinical reports), 11/101 were CD123-negative, and 81/101 were CD123-positive (≥20% of leukemic blasts expressed CD123) of which 50 were high and 31 dim according to clinical pathology reports. Analysis of *inter*patient tagraxofusp-induced priming differences among CD123-positive and CD123-negative groups showed significantly higher delta priming to tagraxofusp in the CD123-positive versus CD123-negative cases (**Fig. 4a, left;** P=0.0042). Of the 81 CD123-positive samples, delta priming values with tagraxofusp monotherapy were only available in 72/81 (89%; 25/72 were dim) as venetoclax-combinations were prioritized where possible when limited by cell number. Further subdividing CD123-positive cases by expression level (high versus dim) showed significant increased delta priming in the CD123-positive cases regardless of expression level (**Fig. 4a, right**), in line with the clinical observation that CD123 expression does not reliably associate with likelihood of clinical response to CD123-directed therapies [23]. To ensure we were not simply observing broad chemosensitivity in CD123-expressing AML cases, we demonstrated that CD123-positive cases showed enhanced sensitivity only to tagraxofusp compared with the other drugs in the panel (**Fig. 4b**, P=0.0063). Interestingly, the two “CD123 negative” cases that primed highest to tagraxofusp monotherapy were later clinically found to have CD123-positive cells detected as evidence of early relapse, suggesting the possibility that DBP could be a means to identify clinically-meaningful AML subpopulations or minimal residual disease (MRD).

**Figure 4.**
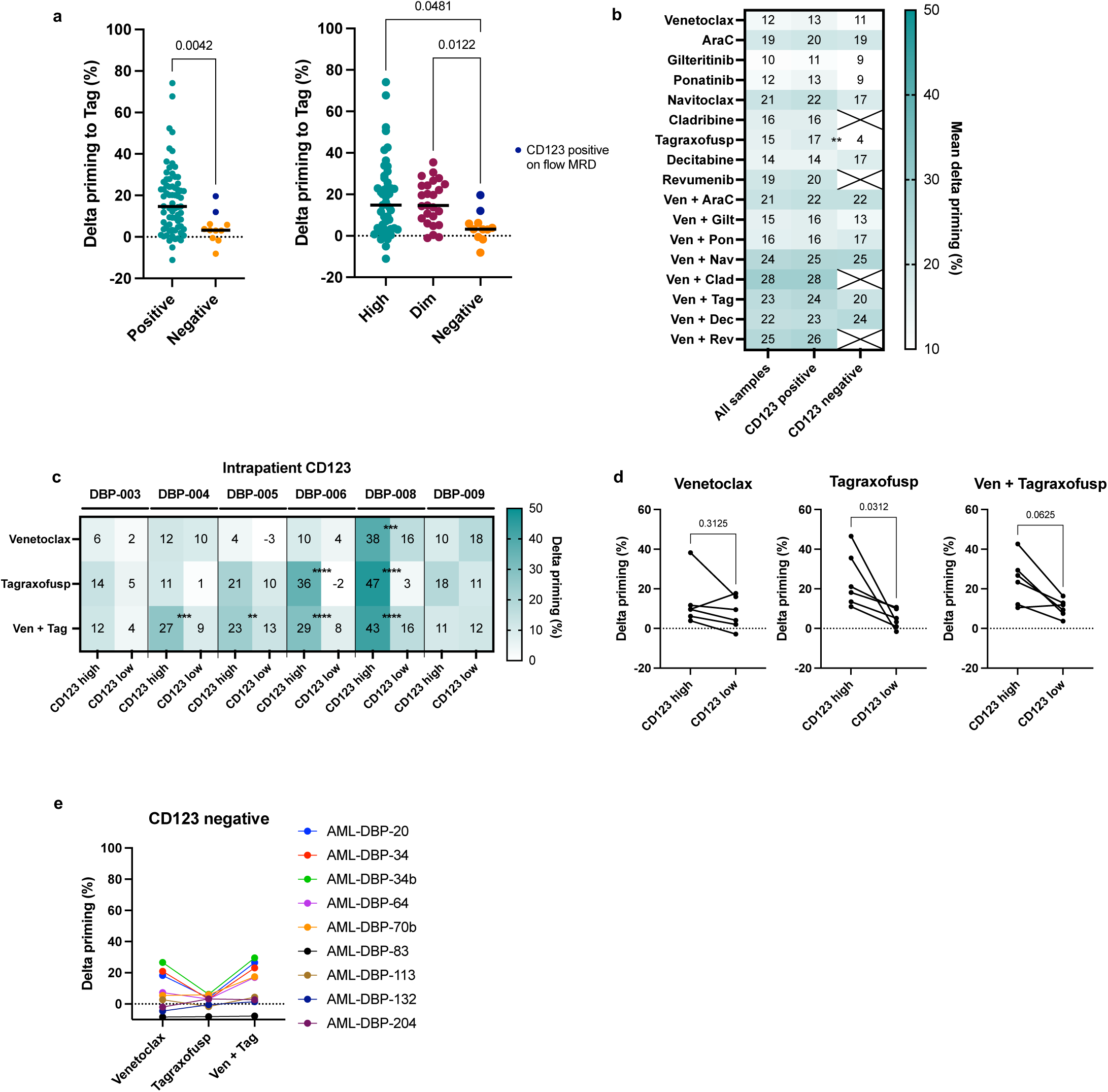
DBP captures tagraxofusp-induced priming changes based on CD123 expression. (a) (left) Comparison of mean delta priming responses to tagraxofusp monotherapy in samples with available CD123 immunophenotyping according to clinical pathology; CD123-positive (n=72) or CD123-negative (n=11) samples (one-tailed Mann-Whitney test). (right) Comparison of mean delta priming to tagraxofusp in CD123 high (n=47), CD123 dim (n=25) and CD123 negative (n=11) samples (one-tailed Mann-Whitney test). (b) Mean delta priming for all treatments for all samples, CD123-positive and CD123-negative samples are compared (one-tailed Mann-Whitney test). (**c**) Myeloblasts (CD33+CD64-CD11b-CD16-) were further gated on CD123 expression and delta priming was compared in high versus low CD123-expressing subpopulations within 6 individual patients treated with venetoclax, azacitidine and tagraxofusp (two-way ANOVA with Tukey’s multiple comparisons; **** p<0.0001, *** p<0.001, ** p<0.01). (**d**) Pairwise comparisons for intrapatient CD123 high versus low subpopulations in response to venetoclax, tagraxofusp, or the combination (Wilcoxon rank test). (**e**) Mean delta priming responses to the indicated treatments in CD123-negative samples.

A major advantage of DBP over other cell death assays is that, as a single cell assessment, it has the ability to observe priming changes within distinct subpopulations, which may be masked by bulk assessment. DBP therefore offers the possibility of identifying drugs that are efficacious against cell subsets of a patient’s myeloblasts that may eventually be responsible for relapse, but additional investigation is warranted with paired diagnosis/relapse samples to confirm. To investigate the extent to which DBP can distinguish heterogeneity of response within patient samples, we determined if CD123-expression levels in this cohort corresponded to apoptotic signaling within myeloblasts in response to tagraxofusp. We identified viably cryopreserved pre-treatment bone marrow samples from six AML patients who were treated with combination venetoclax, azacitidine, and tagraxofusp. Using a CD123 fluorescence minus one (FMO) to determine staining, we gated on CD123-expression in these six CD123-positive cases and assessed delta priming differences in the high (signal above FMO) versus low (signal equal to FMO) expressing myeloblast subpopulations (**Fig. 4c**). We observed significantly higher drug-induced priming to combination venetoclax and tagraxofusp among myeloblasts from 4/6 cases in the CD123-high expressing subpopulations. Delta-priming appears to be driven by tagraxofusp-monotherapy in two cases (DBP-006 and DBP-008), and by venetoclax and tagraxofusp-combination in two others (DBP-004 and DBP-005). Pairwise-comparison revealed no significant differences in venetoclax-induced priming in CD123-high and -low subpopulations (P=0.313), but a significant increase in delta priming was observed in the CD123 high expressing subpopulations in response to tagraxofusp monotherapy (P=0.03) and in combination with venetoclax (P=0.06; **Fig. 4d**). when compared to the CD123-low expressing subpopulations.

Finally, we observed no significant difference in delta priming between CD123-positive and CD123-negative AML samples when treated *ex vivo* with combination venetoclax plus tagraxofusp (**Fig. 4b**). We thus asked whether the addition of venetoclax was dominating apoptotic priming changes in these CD123-negative cells. Indeed, we observed minimal to no contribution from tagraxofusp in CD123-negative samples (**Fig. 4e**). Altogether, these data highlight that DBP can identify drug sensitivity even when they are selectively active only on a subset of the myeloblasts.

### *Ex vivo* drug induced priming predicts clinical outcomes in AML

The ultimate goal of predictive biomarkers is to provide rapid, precise and actionable data to inform individual patient therapy. To test the predictive power of DBP in AML, we first identified patients from our database who received intensive chemotherapy agents and/or targeted therapies included in the fixed-drug panel following DBP. 16 different frontline regimens were administered to the 58 untreated AML patients, and 16 different R/R regimens were administered to the 35 R/R AML patients, which highlights the increasing complexity of modern AML treatment (**Supplemental Table 1**). In newly diagnosed patients who received a 7+3-based (or liposomal encapsulated daunorubicin/cytarabine) after DBP sample collection (N=24), we asked if delta priming to *ex vivo* AraC treatment could predict the achievement of complete remission (CR) or CR with incomplete count recovery (CRi) (**Fig. 5a**). 19 achieved remission and 5 were refractory. With the analyzer blinded to clinical responses until time of data analysis, DBP predicted achievement of CR/CRi in these patients (**Fig. 5a**, P=0.001), providing evidence of its ability to serve as a predictive biomarker (AUC=0.93, P=0.004). These 5 refractory patients had adverse-risk disease without a mutation in *TP53*.

**Figure 5.**
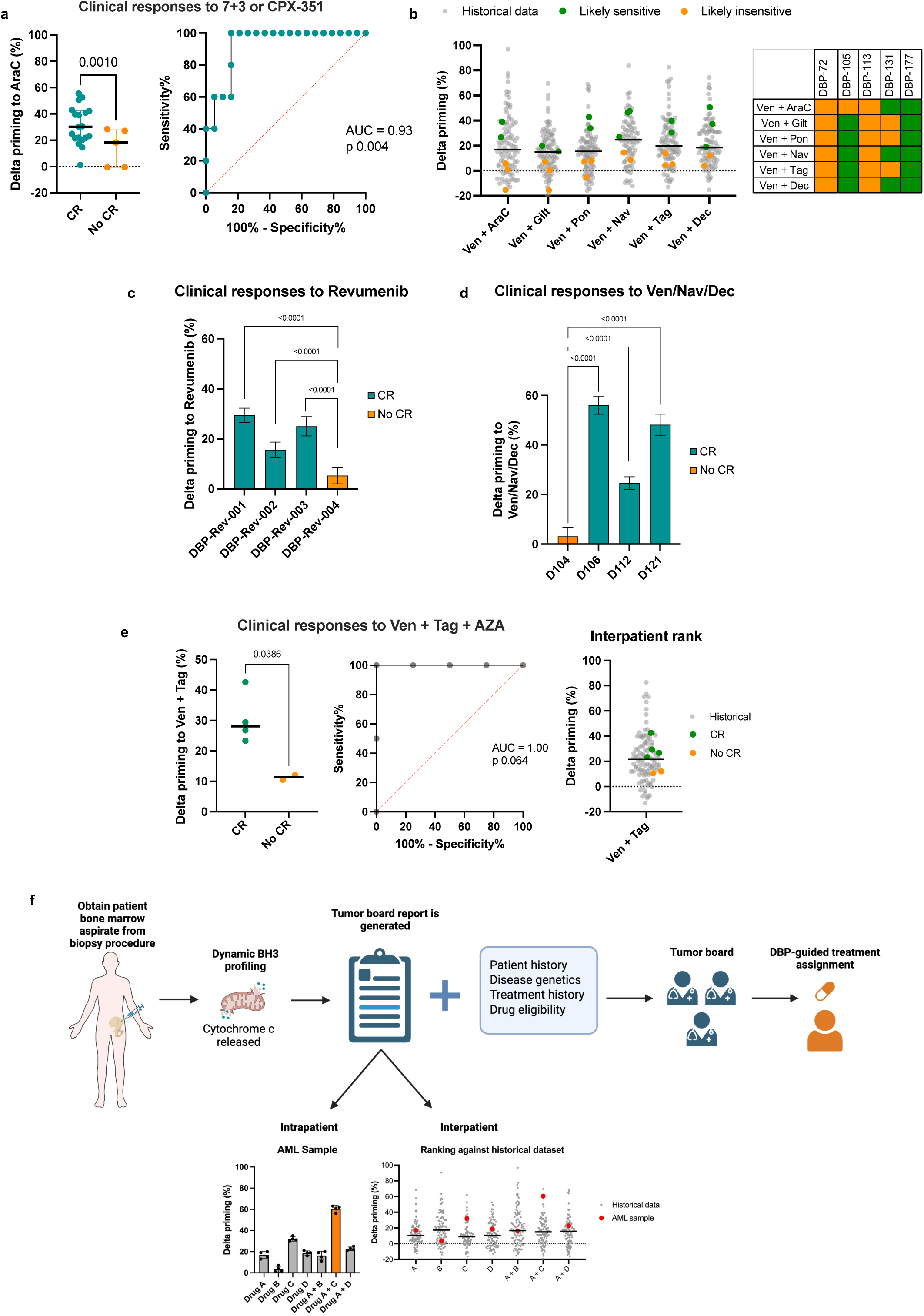
*Ex vivo* DBP in myeloblasts predicts *in vivo* clinical outcomes to intensive chemotherapies and targeted therapy. (**a**) Correlation of delta priming and remission status of patients (N=24) who received 7+3 or CPX-351 after bone marrow was collected for DBP (left) with the corresponding receiver operating characteristic (ROC) curve (right). Black line is at median with interquartile range and p values calculated using one-tailed Mann Whitney test. (**b**) DBP data for venetoclax-based combinations in the 5 patients who did not achieve a CR to 7+3 based therapy. Green = likely sensitive as determined by a delta priming above the historical median delta priming per drug; orange = likely insensitive determined by a delta priming below the historical median per drug. Grey = historical DBP data (n=98 not including samples from the 5 non-responders). Genetics and cytogenetics for each non-responding include: DBP-72 (mutations in *CALR, IDH1, RUNX1, RUNX1,* and *SRSF2*; 47,XY,+8[14]/46,XY[6]); DBP-105 (mutations in *DNMT3A, IDH2, JAK2,* and *EZH2*; 46,XX,del(13)(q12q14)[20]); DBP-113 (mutations in *BCOR, DNMT3A, IDH2, RAD21, SF3B1, STAG2, and TET2*; 46,XX[20]); DBP-131 (mutations in *ASXL1, CBL,* and *EZH2*; 46∼50,XX,inv(3)(q21q26),-7,+9,+13,+22,+22[17]); and DBP-177 (mutations in *DNMT3A, IDH2, RUNX1, SRSF2,* and *FLT3*-ITD; 47,XY,+X[20]). Revumenib and Cladribine were added to the assay after these samples were run. (**c**) Correlation of delta priming and complete remission status of patients who received revumenib from pre-treatment bone marrow samples obtained for DBP (mean ± SD; two-way ANOVA and Dunnett’s multiple comparisons, N=4). (**d**) Correlation of delta priming and remission status of patients who received venetoclax, navitoclax, and decitabine (Ven+Nav+Dec) from pre-treatment bone marrow samples obtained for DBP (mean±SD; two-way ANOVA and Dunnett’s multiple comparisons, N=4). (**e**) Retrospective correlation of *ex vivo* delta priming and CR status of 6 patients who received venetoclax, tagraxofusp, and azacitidine (Ven+Tag+Aza, left; P=0.0386 calculated using one-tailed Mann Whitney test) with the corresponding receiver operating characteristic (ROC) curve (middle). (right) Interpatient ranking of delta priming responses for each sample (4 cases of CR represented by green filled circle and 2 cases of non-responders represented by yellow-orange filled circle) against historical *ex vivo* DBP data with combination venetoclax and tagraxofusp (n=101). (**f**) Proposed model of clinical application of DBP. Fresh bone marrow aspirate obtained from AML patients are subjected to dynamic BH3 profiling and a report is generated from the tumor board. Tumor board reports will contain interpatient and intrapatient data to assess drug-induced apoptotic priming within the sample and against historical data from the AML DBP database. DBP data along with patient history and drug eligibility will be provided to a tumor board for ultimate treatment assignment. CR = complete remission

As DBP predicted futility of 7+3-based therapy in 5 patients, we next asked if there was any evidence of drug sensitivity among the venetoclax-combinations in our assay, including another standard of care frontline option with combination decitabine and venetoclax (**Fig. 5b**). Delta priming values above and below the historical median delta priming per drug condition were termed “likely sensitive” and “likely insensitive,” respectively. Myeloblasts from 2/5 patients showed minimal to no drug sensitivity to any venetoclax-based combination (DBP-72 and DBP-113), and myeloblasts from 1/5 patients showed sensitivity to all combinations tested (DBP-177). Myeloblasts from patient DBP-105 showed minimal to no drug-induced priming to combination venetoclax and AraC, but predicted to be likely sensitive to combination venetoclax and decitabine, providing an example of a patient who may have benefitted from less-intensive therapy.

To validate the predictive ability of DBP, we next identified patients at our institution who had received one of the novel drugs included in our DBP fixed-drug panel and who had viably frozen cells in our tissue bank, but who were not among the 92-patients in the current study. We hypothesized that DBP predicts clinical response to targeted therapies and that its utility was not restricted to intensive chemotherapies. The menin inhibitor revumenib was recently approved for R/R *KMT2A*-rearranged AML [17] with an expected composite CR and complete remission with partial hematological recovery (CRh) rate of 30%. From myeloblasts collected from 4 R/R patients who then went on to receive revumenib on the AUGMENT-101 trial (<u>NCT04065399</u>), we performed DBP to measure revumenib-induced apoptotic priming (**Fig. 5c**). A significantly increased delta priming to revumenib was observed in the responders (N=3) compared to the single non-responder, with all 3 responders achieving a delta priming at least 10-fold higher than the non-responder.

Next, we identified 4 secondary AML patients who received venetoclax, navitoclax and decitabine on clinical trial (NCT05455294). Pre-treatment myeloblasts underwent *ex vivo* drug treatment with the same triplet therapy (**Fig. 5d**). Treatment with combination venetoclax, navitoclax, and decitabine induced significantly higher delta priming in responders (N=3) compared to the non-responder (N=1), with all 3 responders achieving a delta priming at least 20-fold higher than the non-responder.

Finally, to determine if drug-induced priming by tagraxofusp predicted clinical responses to venetoclax, tagraxofusp, and azacitidine (Ven+Tag+Aza), we performed DBP on pre-treatment viably frozen bone marrow mononuclear cells from 6 AML patients who went on to receive this triplet regimen on trial (<u>NCT03113643</u>) [23]. *Ex vivo* drug-induced priming by combination venetoclax and tagraxofusp predicted achievement of complete remission to triplet Ven+Tag+Aza therapy (**Fig. 5e, left**; P=0.0386), with the corresponding receiver operating characteristic (ROC) curve showing excellent biomarker performance (**Fig. 5e**, **middle**; AUC=1.0, P=0.064). DBP provided accurate predictive information across a spectrum of treatments of widely varying mechanism of action. *Inter*patient comparison against our established DBP database (“historical”) shows that patients who achieved CR had delta priming values above the pooled median and those who did not achieve CR had values below, providing evidence to apply ranking.

The AML DBP database continues to accumulate new data, and our goal is to use this database as an index against which future patients can be measured. A proposed model includes presentation of integrated clinical data including patient and prior treatment history, disease genetics and other characteristics (e.g. CD123 status), drug-specific eligibility (e.g. QT interval measurements or organ function), and results from dynamic BH3 profiling to an AML tumor board who can then assign therapies (**Fig. 5f**).

## Discussion

Here, we present the largest dynamic BH3 profiling database on AML patients with prognostic and predictive evidence of utility. We demonstrate DBP can be performed prospectively on primary AML samples with high efficiency, a 16-hour *ex vivo* culture, and a rapid 48-hour turnaround time. We provide proof-of-concept data using *ex vivo* measurements of drug-induced apoptotic priming (likelihood of effectiveness) from a broad range of drugs (as monotherapies and combinations) with real-time response-adaptive therapy. Within our comprehensive AML DBP database, 26 different treatment regimens were administered to 92 AML patients, emphasizing the increasing complexity of AML treatment selection and urgent need for guidance on how to select a patient’s best therapeutic option. No functional biomarkers are currently available in the clinic to provide real-time guidance on therapies that are likely to be efficacious or futile, and the latter would spare patients from unnecessary toxicity. We propose DBP can be positioned in the clinic to prospectively and functionally guide therapeutic decisions in AML, including providing actionable information far before genetic information is reported.

The rapid detection of early apoptosis by DBP provides an advantage over other canonical cell death assays as the myeloblasts are subjected to minimal *ex vivo* culture time which may cause selection, adaptation, or spontaneous cell death. Standard cell death drug response assays often require at least 48–96 hours after drug treatment, and some drugs, like chemotherapy, require time to initiate a cytotoxic effect *ex vivo*. One major advantage of DBP is the ability to detect early changes in pro-apoptotic signalling in less than 24 hours, which commonly precedes detection by other cell death methods including CTG and Annexin V. We show evidence that drug-apoptotic priming by DBP correlates with cytotoxicity. To facilitate rapid clinical decision making, results can be reported to the clinic in as little as 48 hours, even for drugs like some chemotherapeutic agents which are known to require 72 or more hours for a measurable cell death readout.

Pooled DBP results from untreated AML samples reflect both clinical and disease states, and this provides a mechanism behind commonly used clinical AML prognostics. We reveal that the level of drug-induced apoptotic signaling may be the mechanism underlying ELN classification, mPRS status and disease state. Delta priming values above the median corresponded to prolonged PFS in favorable ELN 2022 groups, but not intermediate or adverse risk groups. Although DBP could not be used to distinguish PFS based on the magnitude of delta priming among adverse risk AML patients, we observed a broad range of drug-induced apoptotic priming within myeloblasts from adverse risk ELN AML patients, indicating that DBP in these settings has high potential clinical utility at the individual patient level. With respect to disease genetics, the turnaround time is commonly on the order of 1-2 weeks except for select single genes, and this invariably delays initiation of genetics-guided treatment assignment. Notably, the presence of an actionable mutation insufficiently confers a high likelihood of remission achievement with single-agent therapy, highlighting an opportunity for improvement. *FLT3* inhibition is a prime example where despite the presence of a *FLT3* mutation in a third of all AML cases, only 20-30% of patients will achieve a CR to single-agent gilteritinib [19]. Here, we show that DBP can identify potentially efficacious therapies regardless of disease genetics, with the possibility that the assay may further nominate drug candidates that would otherwise not been considered due to the absence of a genetic biomarker. Lastly, DBP can be used to identify treatment futility in the setting of an actionable mutation, and opportunities where the addition of venetoclax may augment responses to targeted therapies in individual AML patients.

An overall relative reduction in drug-induced priming was observed in R/R compared to untreated AML patients. Our prior publications identified reduced baseline apoptotic priming as a mechanism of acquired resistance that is selected in myeloblasts at relapse [9, 12, 13]. In this study, this phenomenon was similarly observed in the R/R setting, suggesting that myeloblasts in the R/R setting are farther from the apoptotic threshold. A unique benefit to DBP is the ability to identify drug vulnerabilities despite chemoresistant states. Our study was limited to assessments using a fixed-drug panel for the purpose of optimization and potential actionability. However, a larger panel of treatments can theoretically be tested, including novel therapies, when large volume biospecimen collections are available.

*Intra*patient drug-induced priming differences within AML subpopulations may inform clinical responses. As our assay is flow-cytometry based, we successfully identified *intra*patient drug-induced priming differences within distinct AML cell subpopulations. Though bulk AML cell populations indicate *ex vivo* sensitivity to drug, we hypothesize that select cell subpopulations that do not demonstrate similar levels of sensitivity or vulnerability may be left behind after treatment, serving as potential nidus for relapse. As proof-of-concept, we observed significant differences in tagraxofusp-induced priming in CD123-expression (high vs low) among CD123- positive myeloblasts. Additional investigation is warranted with paired diagnostic/relapse samples to confirm this preliminary observation. Knowledge of these potential drug-induced priming differences may inform strategic combinations and next-line therapies and assist in identifying novel drugs that may help to eliminate these potentially-resistant subpopulations.

DBP in AML successfully predicted clinical responses not only to conventional therapy, but also to novel therapy. In the upfront setting, 7+3-based regimens are the conventional treatment choice for patients under the age of 60 years. DBP successfully discriminated untreated AML patients who achieved complete remission to 7+3 or CPX-351 following DBP sample collection and those where the treatment was ultimately futile. DBP could have a role in helping clinicians identify when patients should get intensive versus less-intensive frontline treatments, and potentially identify other therapies that may be effective. We further explored the ability of DBP to predict clinical responses to novel therapy combinations including venetoclax/navitoclax/decitabine and venetoclax/tagraxofusp/azacitidine. With the caveat of low sample number, we observed excellent predictive capability of DBP, affording the exciting prospect that DBP can be positioned in the clinic to identify novel, effective treatment options. DBP data and interpretation will be further refined as more patients are represented in the AML DBP database, including applying ranking to drug-induced apoptotic priming responses and identifying delta priming thresholds at upper or lower quartiles. We propose that DBP data can be used in conjunction with patient clinical data (treatment history, genetics, and drug eligibility), to rapidly provide actionable treatment options for patients (**Fig. 5f**).

In conclusion, we demonstrate the feasibility of performing DBP on myeloblasts from AML patients prospectively. Our results suggest that the dynamic BH3 profiling biomarker may have utility in redefining future treatment for AML. We show the predictive capacity of DBP to identify remission-inducing therapies with both chemotherapeutic and targeted agents, as monotherapy or in combination regimens. In addition to genetic stratification, we propose integration of functional biomarkers as a part of clinical work-up at diagnosis and particularly in the R/R setting where guidance is needed most. We plan to integrate this functional biomarker assay into prospective clinical studies for analytical validation. AML therapy has reached a pivotal turning point as we have been rapidly incorporating targeted therapies in the frontline treatment setting with the goal of preventing disease relapse and prolonging survival. Access and application of prospective functional biomarkers are needed to further accelerate the current state of precision medicine.

## Acknowledgments

Research reported in this publication was supported by philanthropic gifts from the Segal and Counts Families (J.S.G), Helen Gurley Brown Foundation/Dana-Farber Cancer Institute (J.S.G.), the Ted and Eileen Pasquarello Tissue Bank in Hematologic Malignancies, and the National Cancer Institute (NCI) of the National Institutes of Health (NIH) under award numbers CA066996 (R.S., D.J.D., A.L.), K08CA245209 (J.S.G.), and R35CA242427 (A.L.). The content is solely the responsibility of the authors and does not necessarily represent the official views of the NIH or NCI. C.R.R. is supported by an Edward P. Evans Foundation Young Investigator Award, Farmor Foundation, and the Claudia Adams Barr Program in Innovative Basic Cancer Research. Venetoclax, tagraxofusp, ponatinib, and gilteritinib drugs were provided by AbbVie, Stemline, Takeda, and Astellas, respectively.

## Authorship Contributions

E-J.M., J.S.G., and A.L. designed the study; J.S.G., L.T., A.J., K.M., A.A.L., M.R.L., E.S.W., E.C.C, S.S., M.W., R.V., V.V., C.R.R., D.J.D., I.G., and R.M.S. consented the patients and collect biospecimen; E-J.M., L.K., and J.W. processed and performed biomarker assays; E-J.M., G.F., S.H., D.S.N. A.L., and J.S.G. analyzed and interpreted the data and created the figures. All authors contributed to data collection, revision of the manuscript, and approval of the final version.

## Data Sharing

The data generated in this study are available within the article and its supplementary data file. The data generated in this study are otherwise not publicly available as such information could compromise patient consent but may be made available upon reasonable request from the corresponding author. Proposals for access should be sent to the corresponding author.

## Methods

### Patient samples

Between 11/2/2021 and 9/30/2024, fresh primary AML peripheral blood and bone marrow samples were obtained after patients signed an informed consent approved by the Institutional Review Board (Brigham and Women’s Hospital and Dana-Farber Cancer Institute Protocol 01-206) and provided via the Pasquarello Tissue Bank. Patients included had a diagnosis of AML, tissue banking consent (01-206), and planned bone marrow biopsy for routine diagnostics or leukemic blood. All biospecimens were provided with a study ID to the study team. A total of 115 samples were used in this study (101 fresh bone marrow and leukemic blood samples prospectively collected for the AML DBP database, and 15 additional cryopreserved bone marrow mononuclear cell samples obtained from the Pasquarello Heme Tissue bank). Mononuclear cells from 101 fresh primary AML leukapheresis and bone marrow samples were isolated using Vacutainer® CPT tubes (BD Biosciences) and erythrocytes removed by ACK lysis buffer. Viably frozen bone marrow mononuclear cells were obtained from an additional 15 patients from the Pasquarello Tissue Bank for *in vivo* response correlation to *ex vivo* drug-induced priming. Secondary use clinical data for each patient was obtained with IRB approval (DFCI Protocol 17-426) and de-identified clinical data were collected for the AML DBP database. For each sample, a minimum of 8 drug conditions were tested (as venetoclax-based combinations were prioritized), and monotherapies and venetoclax combinations were tested in 96/101 samples based on sufficient cell numbers. In 2024, due to loss of access to navitoclax (pharmaceutical pipeline closed), we made the decision to replace its position on the drug plate with cladribine, which is clinically available. We also added revumenib in 2024 as it was approaching regulatory approval. Among 101 DBP assays, based on timing on when the assay was done with the drug condition changes as described above, up to 17 different drug conditions (9 monotherapy, 8 venetoclax combinations) could be performed pending cell number obtained.

### Patient disease characteristics and outcomes

Patients were deemed evaluable if there was follow-up along with a bone marrow biopsy or persistent leukemic blood (suggestive of no response). ELN 2022 response criteria[3] was then applied based on labs and bone marrow findings. Clinical NGS data using the Rapid Heme Panel (88-gene, 3% VAF sensitivity) was obtained, when possible, at diagnosis or relapse. PFS was defined as time of dynamic BH3 profiling collection until relapse or death (whichever happened first). Demographics and sex were collected and included in Table 1.

### Cell culture and drugs

Primary AML were cultured in StemSpan SFEM II media (StemCell Technologies) containing 1x CD34+ expansion supplement (StemCell Technologies). Cells were kept at a concentration of 1×10^6^ per mL and were incubated in a humidified atmosphere at 37°C with 5% CO_2_. All drugs, with the exception of Tagraxofusp, were dissolved in DMSO (10 mmol/L; Sigma) and stored at −20°C prior to use. Tagraxofusp was dissolved in sterile ddH_2_O and stored at −20°C. All drugs were thawed at room temperature prior to use.

### Dynamic BH3 profiling

BH3 peptide plates were generated using the epMotion 5075L liquid handler (Eppendorf). Peptides (Biosynth) were diluted to 2X concentration in BH3 profiling buffer (150 nM Mannitol, 10 mM HEPES-KOH pH 7.5, 50 mM KCl, 0.02 mM EDTA and EGTA, 0.1% BSA, 5 mM Succinate, and 0.01% Pluronic F-68, Sigma) containing 0.002% digitonin. Dynamic BH3 profiling was carried out on primary AML samples seeded at 100,000 cells per well in a 96-well plate and drug treated using the iDOT One drug dispenser (Dispendix). 16 h later drug was removed and cells were stained with Zombie NIR viability dye (BioLegend) diluted in 1x PBS. Excess dye was removed and cells were stained with an AML staining cocktail diluted in 2% FBS/PBS containing FcR blocking reagent. The antibody staining cocktail included CD3 (BioLegend, RRID:AB-2820225), CD19 (BioLegend, RRID:AB-2820226), CD235a (BioLegend, RRID:AB-2562706), CD33 (BD Biosciences, RRID:AB-400085), CD64 (BioLegend, RRID:AB-2800780), CD11b (BioLegend, RRID:AB-2749870), CD16 (BioLegend), CD45 (BD Biosciences, RRID:AB-2870504), CD123 (BioLegend, RRID:AB-1645547), CD14 (BD Biosciences, RRID:AB-2870488). Cells were resuspended in BH3 profiling buffer and plated into a BH3 profiling peptide plate using the epMotion 5075L liquid handler. 1h later cells were fixed in formalin and neutralized in 3M Tris buffer. Cells were stained with anti-cytochrome c (BioLegend, RRID:AB-2749869) in BD Perm/Wash solution (BD Biosciences) overnight. All flow cytometry was performed on the iQue Screener PLUS (Intellicyt) using the built in ForeCyt software. AML blasts were identified by CD33 positivity (unless pathology confirmed a CD33 negative samples), CD11b and CD16 negativity, and CD45/SSC-low. All subsequent analyses were carried out in Excel (RRID:SCR_016137) or GraphPad Prism (RRID:SCR_002798).

### Annexin V cell death assays

Primary AML samples were used for all annexin V assays. Cells were cultured at a density of 100,000 cells per well of a 96-well plate and treated using the iDOT One drug dispenser (Dispendix). Annexin V Alexa Fluor-488 was diluted to 1μg/mL in 10X Annexin V buffer (100 mM HEPES, 40 mM KCl, 1.4 M NaCl, 25 mM CaCl_2_.2H_2_O, 7.5 mM MgCl_2_, pH 7.4) and added directly to the cells to a final 1X concentration. Annexin V staining was detected by flow cytometry using the iQue Screener PLUS (Intellicyt) and the built in ForeCyt software.

### Statistical methods

For delta priming comparisons, we took the mean of 4-6 BIM 10nM peptide replicates per drug test per sample; thus each patient had up to 17 individual mean drug test results (9 monotherapies and 8 combinations). For pooled delta priming comparisons, the mean of 4-6 BIM peptide replicates per drug per sample was used; thus one sample could provide up to 17 individual data points. For primary analysis (PFS) we determined the mean delta priming values across all drugs per patient and plotted the PFS based on whether that average delta priming value was above or below the pooled median delta priming for the ELN classification.

## References

a1. Bataller, A., H. Kantarjian, A. Bazinet, et al., Outcomes and genetic dynamics of acute myeloid leukemia at first relapse. Haematologica, 2024. 109(11): p. 3543–3556.

2. Burd, A., R.L. Levine, A.S. Ruppert, et al., Precision medicine treatment in acute myeloid leukemia using prospective genomic profiling: feasibility and preliminary efficacy of the Beat AML Master Trial. Nat Med, 2020. 26(12): p. 1852–1858.

3. Dohner, H., A.H. Wei, F.R. Appelbaum, et al., Diagnosis and management of AML in adults: 2022 recommendations from an international expert panel on behalf of the ELN. Blood, 2022. 140(12): p. 1345–1377.

4. Harris, L.N., C.D. Blanke, H.P. Erba, et al., The New NCI Precision Medicine Trials. Clin Cancer Res, 2023. 29(23): p. 4728–4732.

5. Qin, G., J. Dai, S. Chien, et al., Mutation Patterns Predict Drug Sensitivity in Acute Myeloid Leukemia. Clin Cancer Res, 2024. 30(12): p. 2659–2671.

6. Tyner, J.W., C.E. Tognon, D. Bottomly, et al., Functional genomic landscape of acute myeloid leukaemia. Nature, 2018. >562(7728): p. 526–531.

7. Malani, D., A. Kumar, O. Bruck, et al., Implementing a Functional Precision Medicine Tumor Board for Acute Myeloid Leukemia. Cancer Discov, 2022. 12(2): p. 388–401.

8. Kytola, S., I. Vanttinen, T. Ruokoranta, et al., Ex vivo venetoclax sensitivity predicts clinical response in acute myeloid leukemia in the prospective VenEx trial. Blood, 2025. 145(4): p. 409–421.

9. Thanh-Trang Vo, J.R., Ruben Carrasco, Donna Neuberg, Derrick J. Rossi, Richard M. Stone, Daniel J. DeAngelo, Mark G. Frattini, Anthony Letai, Relative Mitochondrial Priming of Myeloblasts and Normal HSCs Determines Chemotherapeutic Success in AML. Cell, 2012. 151(2): p. 344–355.

10. Fraser, C., J. Ryan, and K. Sarosiek, BH3 Profiling: A Functional Assay to Measure Apoptotic Priming and Dependencies. Methods Mol Biol, 2019. 1877: p. 61–76.

11. Deng, J., N. Carlson, K. Takeyama, et al., BH3 profiling identifies three distinct classes of apoptotic blocks to predict response to ABT-737 and conventional chemotherapeutic agents. Cancer Cell, 2007. 12(2): p. 171–85.

12. Bhatt, S., M.S. Pioso, E.A. Olesinski, et al., Reduced Mitochondrial Apoptotic Priming Drives Resistance to BH3 Mimetics in Acute Myeloid Leukemia. Cancer Cell, 2020. 38(6): p. 872–890 e6.

13. Olesinski, E.A., K.S. Bhatia, C. Wang, et al., Acquired Multidrug Resistance in AML Is Caused by Low Apoptotic Priming in Relapsed Myeloblasts. Blood Cancer Discov, 2024. 5(3): p. 180–201.

14. Garcia, J.S., S. Bhatt, G. Fell, et al., Increased mitochondrial apoptotic priming with targeted therapy predicts clinical response to re-induction chemotherapy. Am J Hematol, 2020. 95(3): p. 245–250.

15. Lindsley, R.C., B.G. Mar, E. Mazzola, et al., Acute myeloid leukemia ontogeny is defined by distinct somatic mutations. Blood, 2015. 125(9): p. 1367–76.

16. Dohner, H., C.D. DiNardo, F.R. Appelbaum, et al., Genetic risk classification for adults with AML receiving less-intensive therapies: the 2024 ELN recommendations. Blood, 2024. 144(21): p. 2169–2173.

17. Issa, G.C., I. Aldoss, J. DiPersio, et al., The menin inhibitor revumenib in KMT2A-rearranged or NPM1-mutant leukaemia. Nature, 2023. 615(7954): p. 920–924.

18. Daver, N., A.E. Perl, J. Maly, et al., Venetoclax Plus Gilteritinib for FLT3-Mutated Relapsed/Refractory Acute Myeloid Leukemia. J Clin Oncol, 2022. 40(35): p. 4048– 4059.

19. Perl, A.E., G. Martinelli, J.E. Cortes, et al., Gilteritinib or Chemotherapy for Relapsed or Refractory FLT3-Mutated AML. N Engl J Med, 2019. 381(18): p. 1728–1740.

20. Wei, A.H., P. Panayiotidis, P. Montesinos, et al., 6-month follow-up of VIALE-C demonstrates improved and durable efficacy in patients with untreated AML ineligible for intensive chemotherapy (141/150). Blood Cancer J, 2021. 11(10): p. 163.

21. Kadia, T.M., P.K. Reville, X. Wang, et al., Phase II Study of Venetoclax Added to Cladribine Plus Low-Dose Cytarabine Alternating With 5-Azacitidine in Older Patients With Newly Diagnosed Acute Myeloid Leukemia. J Clin Oncol, 2022. 40(33): p. 3848– 3857.

22. Short, N.J., D. Nguyen, E. Jabbour, et al., Decitabine, venetoclax, and ponatinib for advanced phase chronic myeloid leukaemia and Philadelphia chromosome-positive acute myeloid leukaemia: a single-arm, single-centre phase 2 trial. Lancet Haematol, 2024. 11(11): p. e839–e849.

23. Lane, A.A., J.S. Garcia, E.G. Raulston, et al., Phase 1b trial of tagraxofusp in combination with azacitidine with or without venetoclax in acute myeloid leukemia. Blood Adv, 2024. 8(3): p. 591–602.

24. Chen, E.C., Y. Liu, H. Bell, et al., 4580 Dual Bclxl and BCL2 Inhibition with Navitoclax (NAV), Venetoclax (VEN), and Decitabine (DEC) for Advanced Myeloid Neoplasms (MN): Safety and Biological Activity in a Phase 1 Study, in ASH. 2024.

25. Zeidner, J.F., T.L. Lin, R.L. Welkie, et al., Azacitidine, Venetoclax, and Revumenib for Newly Diagnosed NPM1-Mutated or KMT2A-Rearranged AML. J Clin Oncol, 2025. 43(23): p. 2606–2615.

26. Bataller, A., A. Bazinet, C.D. DiNardo, et al., Prognostic risk signature in patients with acute myeloid leukemia treated with hypomethylating agents and venetoclax. Blood Adv, 2024. 8(4): p. 927–935.

27. Othman, J., N. Potter, A. Ivey, et al., Molecular, clinical, and therapeutic determinants of outcome in NPM1-mutated AML. Blood, 2024. 144(7): p. 714–728.

28. Ivey, A., R.K. Hills, M.A. Simpson, et al., Assessment of Minimal Residual Disease in Standard-Risk AML. N Engl J Med, 2016. 374(5): p. 422–33.

29. Orgueira, A.M., M.P. Encinas, J.A. Diaz Arias, et al. 974 The FLT3-like Gene Expression Signature Predicts Response to Quizartinib in Wild-Type FLT3 Acute Myeloid Leukemia: An Analysis of the Pethema Quiwi Trial. in ASH. 2023.

30. Montesinos, P., R. Rodriguez-Veiga, J.M.B. Burgues, et al. 1512 Final Results of Quiwi: A Double Blinded, Randomized Pethema Trial Comparing Standard Chemotherapy Plus Quizartinib Versus Placebo in Adult Patients with Newly Diagnosed FLT3-ITD Negative AML. in ASH. 2024.

31. Lamble, A.J., L. Eidenschink Brodersen, T.A. Alonzo, et al., CD123 Expression Is Associated With High-Risk Disease Characteristics in Childhood Acute Myeloid Leukemia: A Report From the Children’s Oncology Group. J Clin Oncol, 2022. 40(3): p. 252–261.

32. Pemmaraju, N., A.A. Lane, K.L. Sweet, et al., Tagraxofusp in Blastic Plasmacytoid Dendritic-Cell Neoplasm. N Engl J Med, 2019. 380(17): p. 1628–1637.

