## Supplemental Figures for "Dynamic BH3 profiling predicts clinical outcomes in acute myeloid leukemia"

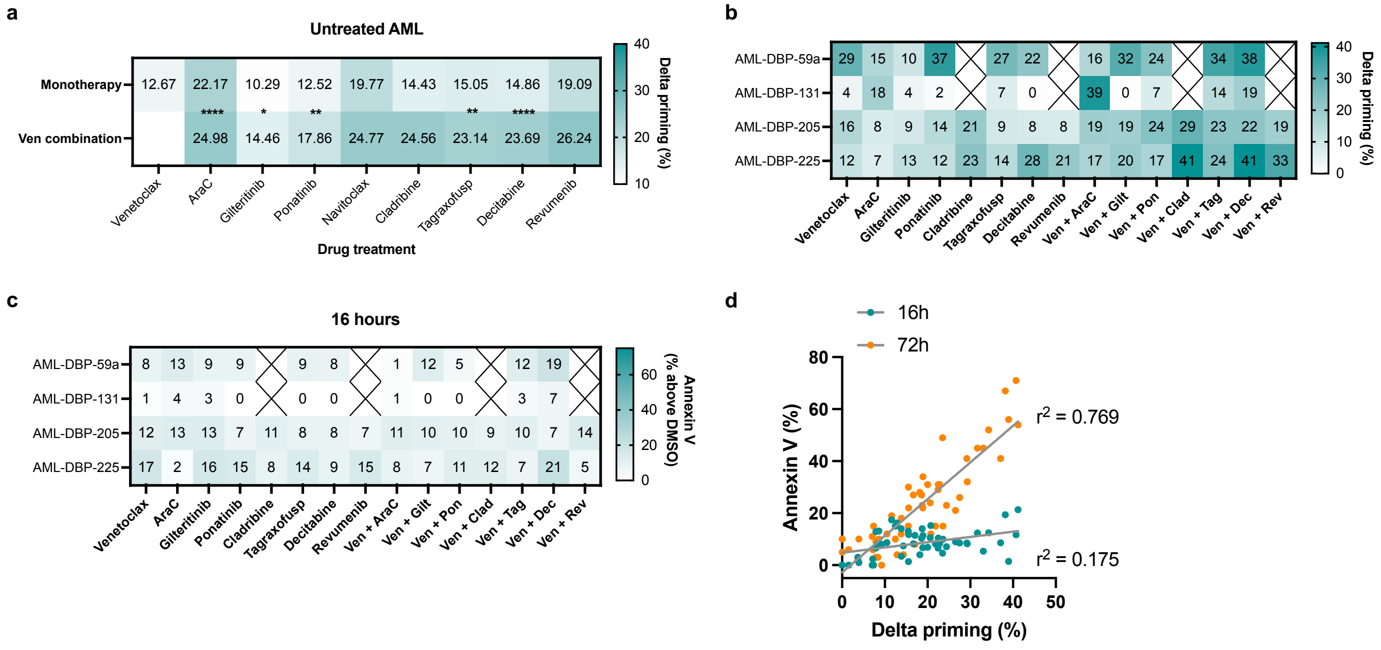


**Supplementary Figure 1 – Delta priming rapidly predicts commitment to cell death in AML.**

(**a**) Mean delta priming for each monotherapy is compared to combination with venetoclax in untreated AML patient samples. Significance determined by two-way ANOVA and Tukey's multiple comparisons test. (**b-d**) Four unique untreated AML patient samples were treated with the indicated drugs. Black cross indicates no data available. (**b**) After 16 hours, BH3 profiling was performed and drug-induced apoptotic priming in myeloblasts was determined. (**c**) Annexin V positivity in myeloblasts was assessed after 16 hours of drug treatment. (**d**) Delta priming (release of cytochrome c) at 16h correlated with cell death at 16h (teal) or 72h (orange). $r^2$ determined by Pearson correlation.

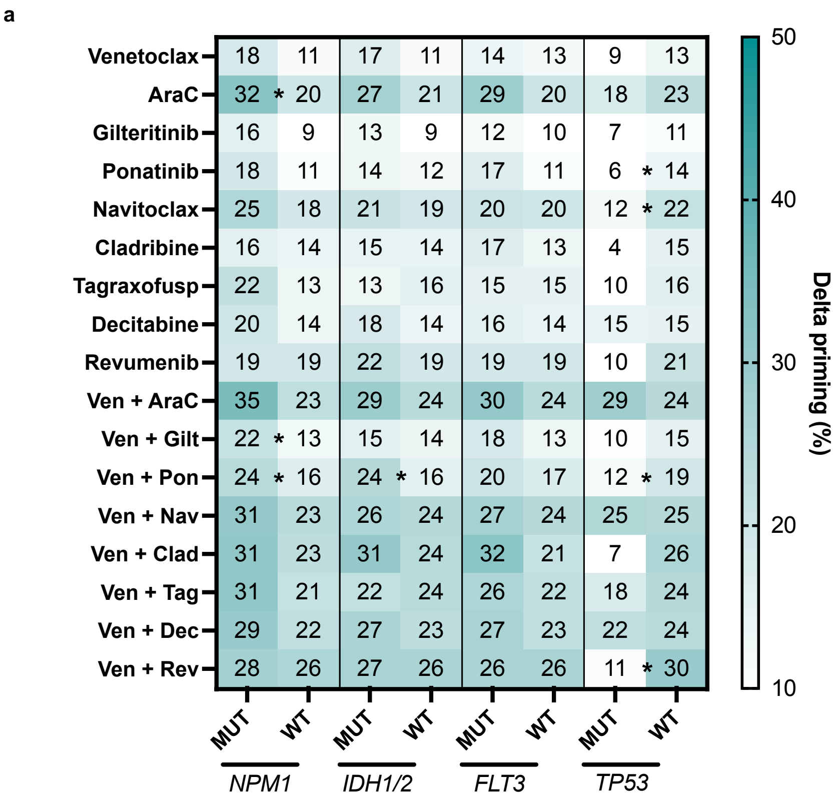


**Supplementary Figure 2 – Delta priming differences observed in AML are not informed by genetics.**

(**a)** Mean delta priming for all treatments for untreated AML samples are compared to *NPM1* mutated (MUT) or wildtype (WT), *IDH1/2* MUT or WT, *FLT3* MUT or WT, and *TP53* MUT or WT. Data represents the mean of 4-6 peptide replicates for drug-induced priming of each drug. Significance determined by unpaired t-test (* = p<0.1).

**Supplementary Table 1**

| Subject Study ID | Age at time of DBP (years) | Disease Histology (WHO5) or ICC 2022 | AML Disease Status at time of collection | Treatment received in clinic after DBP collection | Response status after treatment received by ELN 2022 | Myeloid NGS mutation positive | Mutations Present by NGS | Cytogenetics | CD123 % per Clinical Pathology | CD123 expression? |
|---|---|---|---|---|---|---|---|---|---|---|
| AML-DBP-1 | 77 | Acute myeloid leukaemia, myelodysplasia-related | Refractory | imatinib | no response | Yes | FLT3 D835Y, RAD21, RUNX1, WT1 | 46,XY,t(9;22)(q34;q11.2)[20] | 97 | high |
| AML-DBP-3 | 92 | Acute myeloid leukaemia, myelodysplasia-related | Untreated | Combination venetoclax and azacitidine | no response | Yes | ASXL1, BCOR, GATA2, IKZF1, PHF6, SRSF2, TET2 | 46,XX,add(16)(p13.?2),del(16)(p1?2)[18]/46,XX[1].ish add(16)(MYH11+,CBFB+), del(16)(MYH11-,CBFB+)[3].nuc ish(MYH11x1,CBFBx2)[46/100 | 86 | positive |
| AML-DBP-4 | 66 | AML with defining genetic abnormalities | Relapsed/Refractory | MEC (mitoxantrone, etoposide, cytarabine) salvage chemotherapy | no response | Yes | NPM1, TET2, TET2, FLT3-ITD | 46,XX,t(1;5)(p13;q33),inv(2)(p15-16q3?6)[17]/46,XX[3].ish | 91 | dim |
| AML-DBP-6 | 71 | Acute myeloid leukaemia with other defined genetic alterations | Refractory | Combination venetoclax and decitabine | no response | Yes | DNMT3A, NRAS, NRAS, WT1, WT1, FLT3-ITD | 46,XX[20] | 99 | N/A |
| AML-DBP-7 | 89 | AML with monocytic differentiation | Untreated | Combination venetoclax and azacitidine | CRi | Yes | ASXL1, PTPN11, PTNP11, RUNX1, FLT3-ITD | 46,XX[20] | 97 | small subset |
| AML-DBP-9b | 82 | Acute myeloid leukaemia, myelodysplasia-related | Refractory | low dose CPX-351/venetoclax on clinical trial | no response | Yes | PTPN11, SF3B1, TP53, TP53 | 47,XY,add(5)(q11.2),+19,?r(22)(p11.2q1?2)[8]/48,idem ,+Y[5]/48,idem,+Y,add(7) (q22)[2]/46,XY[5] | 46 | variable |
| AML-DBP-11 | 85 | Acute myeloid leukaemia, myelodysplasia-related | Relapsed | gilteritinib | no response | Yes | DNMT3A, DNMT3A, NPM1, NRAS, TET2, FLT3-ITD | n/a | 91 | dim |
| AML-DBP-14 | 26 | Acute myeloid leukaemia, myelodysplasia-related | Refractory | clofarabine/cytarabine | no response | Yes | BCOR, GATA2, RUNX1 | 48,XY,add(7)(q36),+8,+21[10] | 93 | high |
| AML-DBP-15 | 78 | Acute myeloid leukemia with myelolmonocytic differentiation | Refractory | no treatment received (supportive care only) | no response | No | none detected | 45,X,-Y,t(11;17)(q23;q?12)[20] | n/a | n/a |
| AML-DBP-18 (1) | 76 | Acute myeloid leukaemia, myelodysplasia-related | Refractory | Combination venetoclax and decitabine | no response | Yes | DNMT3A, RUNX1, SF3B1, TET2, TET2, FLT3-ITD | n/a | 92 | high |
| AML-DBP-19 | 79 | Acute myeloid leukaemia with other defined genetic alterations | Refractory | Combination venetoclax and decitabine | no response | Yes | DNMT3A, TP53, TP53 | 45~46,XY,del(3)(q21q28)[3],del(5)(q15q32),-7,+8,psu dic(12;20)(p11.2;q13.3), +r[3][cp8]/46,XY[10] | 19 | dim to negative |

| | | | | | | | | | | |
|---|---|---|---|---|---|---|---|---|---|---|
| AML-DBP-20 | 61 | Acute myeloid leukaemia with other defined genetic alterations | Refractory | CA-4948 on clinical trial | no response | Yes | CEPBA, TET2, TET2, WT1, WT1 | 46,XY,del(9)(q13q32),add(16)(p13.3),del(19)(p13.1)[3]/ 46,XY[17] | 5 | dim to negative |
| AML-DBP-23 | 69 | Acute myeloid leukaemia, myelodysplasia-related | Relapsed | aza/ven/tagraxofusp on clinical trial | no response | Yes | ASXL1, PHF6, RUNX1, U2AF1 | 46,XY[20] | 29 | dim |
| AML-DBP-27 | 45 | AML with defining genetic abnormalities | Refractory | no treatment received (supportive care only) | no response | Yes | ASXL1, BCOR, GATA2, NRAS, SF3B1, SF3B1 | 46,XY,inv(3)(q21q26.2)[20] | 97% | positive |
| AML-DBP-34 (1) | 27 | Acute myeloid leukaemia with other defined genetic alterations | Relapsed | clofarabine/cytarabine | CR flow MRD positive | Yes | NRAS, PHF6 | 46,XY,add(3)(p13),add(4)(p16),add(4)(q31),del(5)(q2?2q3?2),-9,-11,-15,-16,add(1 7)(q11.2),+4mar[cp10]/46,XY[5] | n/a | negative |
| AML-DBP-34 (2) (biological duplicate) | 27 | Acute myeloid leukaemia with other defined genetic alterations | Relapsed | clofarabine/cytarabine | CR flow MRD positive | Yes | NRAS, PHF6 | 46,XY,add(3)(p13),add(4)(p16),add(4)(q31),del(5)(q2?2q3?2),-9,-11,-15,-16,add(1 7)(q11.2),+4mar[cp10]/46,XY[5] | n/a | n/a |
| AML-DBP-36 | 76 | Acute myeloid leukaemia, myelodysplasia-related | Relapsed | Combination venetoclax and decitabine | no response | Yes | DNMT3A, DNTM3A, U2AF1 | 46,XY[20] | 90 | dim |
| AML-DBP-37b | 73 | Acute myeloid leukaemia, myelodysplasia-related | Untreated | Combination venetoclax and azacitidine | non-evaluable for response | Yes | ASLX1, ASXL1, BCOR, CEBPA, ETV6, KRAS, PTPN11, SETBP1, SRSF2, STAG2 | 46,XY,i(17)(q10)[20] | 19 | variable |
| AML-DBP-38 | 67 | Acute myeloid leukaemia, myelodysplasia-related | Refractory | no treatment received (supportive care only) | non-evaluable for response | Yes | TP53 | 42-47,XX,add(1)(q32),-5,-9,del(9)(p13),del(12)(p11.2),add(14)(p11.2)[2], -16[4],add(16)(p13.3)[7],-19,-21[4],i(21)(q10)[4],+3-6mar[cp10]/46,XX[2] | 26 | dim, partial |
| AML-DBP-39a | 35 | AML with defining genetic abnormalities | Untreated | 7+3 + Gemtuzumab | CR flow MRD negative | Yes | KIT, KIT | 46,XY,inv(16)(p13.1q22) [20]; FISH positive for CBFB rearrangement. | 93 | positive |
| AML-DBP-42a | 69 | Acute myeloid leukaemia with other defined genetic alterations, post cytotoxic chemotherapy | Untreated | aza/ven/tagraxofusp on clinical trial | non-evaluable for response | Yes | TP53, TP53 | 57~59,XX,+1,add(1)(p13),+3,del(3)(q21q26.1),+5,del(5 )(q23q35))x2,+6,+6,del(6) (q23q25)x2,+8,+8,inv(9)(p12q13)?c,+11,add(11)(p1?5) ,+15,+18,+19,-21,+mar1, +2-3mar[cp20]. nuc ish(RP11-669C7/RP11-637O11,RP11-82C9,RP11-362K14)x3[43/100], (NUP98x2)[100] | 90 | positive |
| AML-DBP-49 | 77 | MDS/AML (10% blasts and TP53 mut) | Untreated | decitabine | CR | Yes | ASXL1, BCOR, DNMT3A, STAG2, TP53, U2AF1 | 46,XY,del(20)(q11.2q13.1)[cp6]/46,XY[14] | n/a | negative |
| AML-DBP-51 (1) | 53 | Acute myeloid leukaemia with other defined genetic alterations | Relapsed/ Refractory | MEC (mitoxantrone, etoposide, cytarabine) salvage chemotherapy | no response | Yes | CEPBA, FLT3-ITD, NRAS, PTPN11, TET2, TET2, TET2, WT1, WT1 | 46,XX[20] | 98 | positive |
| AML-DBP-53 | 73 | Acute myeloid leukaemia, myelodysplasia-related, post-MDS | Previously treated for MN, untreated AML | CPX-351 | no response | Yes | BCOR, DNMT3A, TET2,U2AF1 | 46,XY,del(2)(q2?3q3?7),add(5)(q2?2),der(14)t(1;14)(q 21;p11.2),del(20) (q11.2q13.3)[cp7]/46,XY[8].nuc ish(D5S723/D5S721x2,EGR1x1)[81/100] | 78 | positive |

| | | | | | | | | | | |
|---|---|---|---|---|---|---|---|---|---|---|
| AML-DBP-56 | 73 | B/Myeloid Mixed Phenotype Acute Leukemia | Relapsed | Combination venetoclax and decitabine | no response | Yes | U2AF1 | 46,XY,del(5)(q15q33)[1]/46,XY[19].nuc ish(D5S723/D5S721x2,EGR1x1)[12/100] | 67 | positive |
| AML-DBP-58 | 76 | AML with defining genetic abnormalities | Untreated | ven/aza/ivosidenib on clinical trial | CR flow MRD negative | Yes | DNMT3A, DNMT3A, FLT3 D835Y, IDH1, NPM1, PTPN11 | 46,XX[20] | 94 | variable |
| AML-DBP-59 (1) | 73 | Acute myeloid leukaemia, myelodysplasia-related | Untreated | no treatment received (supportive care only) | non-evaluable for response | No | none detected | 42~43,XX,der(2)t(2;3)(p23;p21),der(3)add(3)(p21)add(3)(q21),-4, del(5)(q15q35),-7,i(8)(q10),del(9)(q22q34),del(10)(p11.2),-12,-13,-14[14], -15[17],-16,-17,-18[3],add(22)(q11.2),+3~7mar[cp19]/46,XX[1] | 93 | dim |
| AML-DBP-59 (2) (biological duplicate) | 73 | Acute myeloid leukaemia, myelodysplasia-related | Untreated | no treatment received (supportive care only) | non-evaluable for response | No | none detected | 42~43,XX,der(2)t(2;3)(p23;p21),der(3)add(3)(p21)add(3)(q21),-4, del(5)(q15q35),-7,i(8)(q10),del(9)(q22q34),del(10)(p11.2),-12,-13,-14[14], -15[17],-16,-17,-18[3],add(22)(q11.2),+3~7mar[cp19]/46,XX[1] | 93 | dim |
| AML-DBP-60 | 55 | AML with defining genetic abnormalities | Untreated | 7+3+venetoclax on clinical trial | CR flow MRD negative | Yes | IDH2, NPM1, TET2 | 46,XY[20] | n/a | negative |
| AML-DBP-63 (1) | 78 | Acute myeloid leukaemia, myelodysplasia-related | Previously treated for MN, untreated AML | no treatment received (supportive care only) | non-evaluable for response | Yes | ASXL1, BCOR, EZH2, NRAS, RUNX1, RUNX1, STAG2, TET2 | 47,XY,+8[cp17]/47,idem,del(16)(q?12-13q?24)[2]/46,XY[1] | n/a | n/a |
| AML-DBP-63 (2) (biological duplicate) | 78 | Acute myeloid leukaemia, myelodysplasia-related | Previously treated for MN, untreated AML | no treatment received (supportive care only) | non-evaluable for response | Yes | ASXL1, BCOR, EZH2, NRAS, RUNX1, RUNX1, STAG2, TET2 | 47,XY,+8[cp17]/47,idem,del(16)(q?12-13q?24)[2]/46,XY[1] | n/a | n/a |
| AML-DBP-64 | 60 | Acute myeloid leukaemia, myelodysplasia-related | Untreated | 7+3+venetoclax on clinical trial | CR flow MRD negative | Yes | ASXL1, DNMT3A, IDH2 | 46,XX[20] | 4 | very dim |
| AML-DBP-65 | 89 | Acute myeloid leukaemia, myelodysplasia-related | Refractory | gilteritinib | no response | Yes | ASXL1, CBL, EZH2, FLT3 D835_M837, RUNX1, RUNX1, TET2 | 46,XY[20] | n/a | n/a |
| AML-DBP-66 | 90 | Acute myeloid leukaemia, myelodysplasia-related | Untreated | Combination venetoclax and azacitidine | CR flow MRD positive | Yes | ASXL1, TET2, TET2 | 46,XX,t(4;12)(q12;p13)[18]/46,XX[2] | 78 | positive |
| AML-DBP-67 | 79 | Acute myeloid leukaemia, myelodysplasia-related | Previously treated for MN, untreated AML | ven/aza/ivosidenib on clinical trial | CR flow MRD negative | Yes | ASXL1, IDH1, RUNX1, SRSF2, TET2 | 46,XY,?del(12)(p13p13)[5]/46,XY[cp15] | 73 | positive |
| AML-DBP-47b | 65 | Acute myeloid leukaemia, myelodysplasia-related, post-MPN | Relapsed/ Refractory | Combination venetoclax and decitabine | no response | Yes | GNB1, KRAS, SRSF2, STAG2, TET2 | 47,XX,+8[20] | 77 | dim |
| AML-DBP-68 | 82 | Acute myeloid leukaemia, myelodysplasia-related | Relapsed | enasidenib | no response | Yes | DNMT3A, IDH2, NF1, NF1, PHF6, PTPN11, RUNX1 | 47,XY,-7,+r(7)(p?q?),+mar1[cp15]/47,XY,+mar1[5] | 54 | dim |

| | | | | | | | | | | |
|---|---|---|---|---|---|---|---|---|---|---|
| AML-DBP-70 (1) | 76 | Acute myeloid leukaemia with other defined genetic alterations | Untreated | Combination venetoclax and azacitidine | MLFS | Yes | KRAS, NRAS, PTPN11, TP53 | 45~49,XX,add(5)(q11.2),-7,+8,add(12)(p11.2),add(17)(p11.2),-18,+1~4r[cp20] | 26 | dim to negative |
| AML-DBP-70 (2) (biological duplicate) | 76 | Acute myeloid leukaemia with other defined genetic alterations | Untreated | Combination venetoclax and azacitidine | MLFS | Yes | KRAS, NRAS, PTPN11, TP53 | 45~49,XX,add(5)(q11.2),-7,+8,add(12)(p11.2),add(17)(p11.2),-18,+1~4r[cp20] | 26 | dim to negative |
| AML-DBP-72 | 66 | Acute myeloid leukaemia, myelodysplasia-related | Untreated | 7+3 | no response | Yes | CALR, IDH1, RUNX1, RUNX1, SRSF2 | 47,XY,+8[14]/46,XY[6] | 55 | dim |
| AML-DBP-51 (2) (biological duplicate) | 53 | Acute myeloid leukaemia with other defined genetic alterations | Relapsed/ Refractory | HIDAC (high dose cytarabine) | no response | Yes | CEPBA, FLT3-ITD, NRAS, PTPN11, TET2, TET2, TET2, WT1, WT1 | 46,XX[20] | 85 | positive |
| AML-DBP-79 | 60 | Acute myeloid leukaemia, myelodysplasia-related | Relapsed | Combination venetoclax and decitabine | no response | Yes | DNMT3A, TP53 | n/a | 16 | variable |
| AML-DBP-82 | 75 | Acute myeloid leukaemia, myelodysplasia-related, post-MDS | Untreated | ven/aza/ivosidenib on clinical trial | CRi flow MRD negative | Yes | IDH1, RUNX1, SRSF2 | 92<4n>,XXYY,del(5)(q12q3?3),+13,+13,-21,-21[cp3]/46,XY[17] | 96 | positive |
| AML-DBP-83 | 52 | Acute myeloid leukaemia, myelodysplasia-related | Untreated | Combination venetoclax and azacitidine | CR flow MRD negative | Yes | TP53 | 44,XX,del(1)(q21),-3,der(5)t(1;5)(q21;q22),add(7)(p13),+8,add(8)(p11.2),-11, -12,add(18)(q11.2),add(20)(p13)[20] | 1 | dim to negative |
| AML-DBP-91 | 84 | Acute myeloid leukaemia with other defined genetic alterations | Untreated | Combination venetoclax and azacitidine | no response | Yes | NPM1, TET2, TET2, FLT3-ITD | 46,XX[20] | 98 | dim |
| AML-DBP-93 | 63 | Acute myeloid leukaemia, myelodysplasia-related, post-MDS | Previously treated for MN, untreated AML | no treatment received (supportive care only) | no response | Yes | ATM | n/a | 96 | dim |
| AML-DBP-96 | 69 | Acute myeloid leukaemia with other defined genetic alterations | Untreated | 7+3+midostaurin | CR | Yes | FLT3-ITD, FLT3 TKD (D835Y), FLT3 (N587_E598dup), FLT3 (N676K), | 46,XY[13] | 91 | dim |
| AML-DBP-100 | 77 | Acute myeloid leukaemia, myelodysplasia-related, with germline predisposition | Untreated | ven/aza/ivosidenib on clinical trial | CR flow MRD negative | Yes | DDX41, U2AF1, PTPN11, IDH1, BCOR | 46,XY[20] | 92 | dim |
| AML-DBP-105 | 68 | Acute myeloid leukemia with myelolmonocytic differentiation | Untreated | 7+3 | no response | Yes | DNMT3A, IDH2, JAK2, EZH2 | 46,XX,del(13)(q12q14)[20] | 55 | dim |
| AML-DBP-107 | 87 | Acute myeloid leukaemia, myelodysplasia-related | Untreated | azacitidine | non-evaluable for response | Yes | NF1 and TP53 | 47,XX,-3,der(5)add(p15)add(q11.2),inv(7)(q22q32),+8,add(11)(q23), del(11)(q23),del(12)(q10),+13,add(13)(p10)x2,del(13)(?q22q34),-17,del(18)(q21), +mar[19]/46,XX[1] | 37 | subset |
| AML-DBP-108 | 69 | Acute myeloid leukaemia, myelodysplasia-related, post-MDS | Relapsed | MEC (mitoxantrone, etoposide, cytarabine) salvage chemotherapy | no response | Yes | ASXL1, RUNX1 | 47,XY,del(5)(q22q35),+21[3]//47,XXY[17] | 77 | subset |

| | | | | | | | | | | |
|---|---|---|---|---|---|---|---|---|---|---|
| AML-DBP-113 | 68 | Acute myeloid leukaemia, myelodysplasia-related | Untreated | CPX-351 | no response | Yes | BCOR, DNMT3A, IDH2, RAD21, SF3B1, STAG2, TET2 | 46,XX[20] | 3 | dim |
| AML-DBP-114 | 77 | Acute myeloid leukaemia with other defined genetic alterations | Refractory | Combination venetoclax and azacitidine | no response | Yes | DNMT3A, TP53, TP53 | no metaphases | 72 | variable |
| AML-DBP-116 (1) | 75 | Acute myeloid leukaemia with other defined genetic alterations | Untreated | 7+3+midostaurin | CR flow MRD negative | Yes | DNMT3A, IDH1, NPM1, PRPF8, WT1, FLT3-ITD | 46,X,t(X;13)(p11.2;q34)[7]/46,XX[13].nuc ish(KMT2Ax2)[100] | 41 | dim, partial |
| AML-DBP-116 (2) (biological duplicate) | 75 | Acute myeloid leukaemia with other defined genetic alterations | Untreated | 7+3+midostaurin | CR flow MRD negative | Yes | DNMT3A, IDH1, NPM1, PRPF8, WT1, FLT3-ITD | 46,X,t(X;13)(p11.2;q34)[7]/46,XX[13].nuc ish(KMT2Ax2)[100] | 41 | dim, partial |
| AML-DBP-121 | 69 | Acute myeloid leukaemia, myelodysplasia-related | Untreated | 7+3+midostaurin | MLFS | Yes | ASXL1, NRAS, RUNX1, RUNX1, SRSF2, STAG2, STAG2, FLT3-ITD | 46,XY,del(11)(q2?2q25)[2]/46,XY[18].nuc ish(MLLx2)[100],(PML,RARA)x2[100] | 15 | dim, subset |
| AML-DBP-124 | 41 | Acute myeloid leukaemia with other defined genetic alterations | Refractory | revumenib on clinical trial | CR flow MRD negative | Yes | CREBBP | 46,XY,t(6;11)(q27;q23)[7]/92<4n>,XXYY,t(6;11)x2[cp4] /46,XY[6] | 77% | High |
| AML-DBP-125 (1) | 69 | Acute myeloid leukemia with monocytic differentiation, post cytotoxic chemotherapy | Relapsed/ Refractory | LY3214996 on clinical trial | no response | Yes | ASXL1, CBL, CBL, CBL, DNMT3A, IKZF1, NRAS, PTPN11, RUNX1, RUNX1, SRSF2 | 46,XY,+1,der(1;7)(q10;p10),t(7;17)(p22;q11.2)[2]/46,X Y[18] | 58% | variable |
| AML-DBP-126 | 92 | Acute myeloid leukaemia, myelodysplasia-related | Untreated | Combination venetoclax and decitabine | non-evaluable for response | Yes | ASXL1, JAK2, RUNX1, TET2, TET2, TP53 | 58- 63,XX,+X,+2,add(2)(p13),+6,+6,+6,t(6;14)(p23;q32),+8,- 9, - 12,+13,+14,+15,+19,+20,add(21)(q22),+22,add(22)(q1 3)+2-8mar[cp20] | 28 | variable,dim |
| AML-DBP-129 | 70 | Acute myeloid leukaemia, myelodysplasia-related | Untreated | no treatment received (supportive care only) | non-evaluable for response | Yes | ASXL1, BCOR, NRAS, PHF6, TET2 | 46,XX,+1,der(1;21)(q10;q10)[20] | 60 | subset, dim |
| AML-DBP-130 | 51 | Acute myeloid leukemia with monocytic differentiation | Untreated | 7+3 | CR flow MRD negative | Yes | ETV6::FLT3 gene fusion | 49,XX,+8,+11,+12[20] | 79 | variable |
| AML-DBP-131 | 78 | Acute myeloid leukaemia, myelodysplasia-related | Untreated | CPX-351 | no response | Yes | ASXL1, CBL, EZH2 | 46~50,XX,inv(3)(q21q26),- 7,+9,+13,+22,+22[17][cp20].nuc ish(RP11-669C7/RP11-637O11,RP11-82C9,RP11- 362K14x2),(RP11-669C7/RP11-637O11 sep RP11-82C9,RP11-362K14x1)[40/100] | 97 | high |
| AML-DBP-125 (2) (biological duplicate) | 69 | Acute myeloid leukemia with monocytic differentiation, post cytotoxic chemotherapy | Relapsed/ Refractory | BXCL701 on clinical trial | no response | Yes | ASXL1, CBL, CBL, CBL, DNMT3A, GATA2, IKZF1, NRAS, PTPN11, RUNX1, RUNX1, | 46,XY[cp20] | 58 | variable |

| | | | | | | | | | | |
|---|---|---|---|---|---|---|---|---|---|---|
| AML-DBP-132 | 59 | Acute myeloid leukaemia with other defined genetic alterations | Untreated | aza/magrolimab on clinical trial | no response | Yes | TP53, TP53 | 47-51,XY,+5[9],del(5)(q13q33),+6,add(11)(p13)[4],psudic(11;5)(p13;p15)[5],der(13;21)(q10;q10)[2],t(19;21)(p13;p11.2),+21[6], add(21)(p11.2)[4],add(21)(p11.2)[3],+1-3 mar[cp18]/45,XY,-4,del(7)(q22q32), del(9)(q22q34),+r[2].nuc ish(D5S723/D5S721x3-6,EGR1x2-4)[20/100]/ (D5S723/D5S721,EGR1)x3-4[11/100],(D7Z1x2,D7S486x1)[57/100]/(D7Z1,D7S486)x3-4[18 /100] | 3 | dim to neg |
| AML-DBP-136 | 78 | Acute myeloid leukemia with myelolmonocytic differentiation | Untreated | HIDAC (high dose cytarabine) | non-evaluable for response | Yes | NPM1, IDH1, FLT3 TKD Y572C, FLT3-ITD | 46,XY[20] | 98 | high |
| AML-DBP-143 | 51 | AML with defining genetic abnormalities | Untreated | 7+3+midostaurin | CR flow MRD negative | Yes | IDH2, NPM1, FLT3-ITD | 46,XX[20].nuc ish(KMT2Ax2)[100] | 48 | dim |
| AML-DBP-145 | 76 | Acute myeloid leukaemia, myelodysplasia-related | Relapsed | revumenib on clinical trial | CR flow MRD negative | Yes | BRCC3, CREBBP, FLT3 D835Y, IDH2, NPM1, SETD2, SRSF2 and TET2 | 46,XY,t(18;21)(q11.2;q22)[17]/46,XY[3] | 96 | positive |
| AML-DBP-148 | 45 | Acute myeloid leukaemia with other defined genetic alterations | Relapsed | CA-4948 on clinical trial | CRi flow MRD negative | Yes | NRAS, WT1, WT1, FLT3-ITD | 46,XY,t(6;9)(p23;q34)[20] | 82 | positive |
| AML-DBP-149 | 66 | AML with defining genetic abnormalities | Untreated | 7+3 | CR flow MRD negative | Yes | CEBPA, CEBPA, EP300 | 46,XX,?del(20)(q13.1q13.3)[13]/46,XX[7].nuc ish(D20S108x2)[100] | 52 | partial |
| AML-DBP-153 | 76 | MDS with excess blasts | untreated | no treatment received (supportive care only) | no response | Yes | BCOR, BCORL1, DNMT3A, EP300, PTPN11, RUNX1, TET2, U2AF1 | 46,XY[20] | 52 | dim |
| AML-DBP-176 | 79 | Acute myeloid leukaemia, myelodysplasia-related | Previously treated for MN, untreated AML | Combination venetoclax and decitabine | no response | Yes | DNMT3A, TET2 TET2, and U2AF1 | 46,XY,del(20)(q11.2q13.3)[5]/47,idem,+20[4] | 1 | dim, partial |
| AML-DBP-177 | 62 | Acute myeloid leukaemia, myelodysplasia-related | Untreated | 7+3+midostaurin | no response | Yes | DNMT3A, IDH2, RUNX1, SRSF2, FLT3-ITD | 47,XY,+X[20] | 73 | partial |
| AML-DBP-178 | 79 | Acute myeloid leukaemia with other defined genetic alterations | Relapsed | no treatment received (supportive care only) | non-evaluable for response | Yes | NRAS and TET2 | 46,XY,+1,der(1;14)(q10;q10)[20] | 86 | partial |
| AML-DBP-182 | 69 | Acute myeloid leukemia with myelolmonocytic differentiation | Relapsed/ Refractory | no treatment received (supportive care only) | non-evaluable for response | Yes | ASXL1, EZH2, EZH2, NF1, NF1 and RUNX1 | 46,XY,t(12;18)(p11.2;q11.2)[7]/46,XY,del(11)(?p15?p13)[cp4]/ 46,XY,t(1;17)(p34;q11.2)[3]/46,XY,t(7;11)(p13;q23)[3]/46,XY[2]//46,XX[1] .nuc ish(MLLx2)[100] | 1 | difficult to assess |
| AML-DBP-186 | 58 | Acute myeloid leukaemia, myelodysplasia-related | Untreated | aza/ven/IMGN632 on clinical trial | CR flow MRD negative | Yes | CEBPA, RAD21, and TET2 | 47,XY,+4[17]/47,idem,del(11)(q13q23)[3] | 39 | variable |

| | | | | | | | | | | |
|---|---|---|---|---|---|---|---|---|---|---|
| AML-DBP-187 | 87 | Acute myeloid leukemia with myelolmonocytic differentiation | Untreated | no treatment received (supportive care only) | non-evaluable for response | Yes | ASXL1 | 46,XY,t(9;11)(p21;q23)[cp18]/46,XY[2] | 63 | dim, partial |
| AML-DBP-190 | 69 | AML with defining genetic abnormalities | Relapsed | ven/aza/ivosidenib on clinical trial | CR flow MRD negative | Yes | ATRX, BRCC3, DNMT3A, IDH1, KRAS, NPM1, U2AF1 | 46,XY[20] | 88 | majority |
| AML-DBP-191 | 81 | MDS/AML (10% blasts and MDS-related gene mutation) | Previously treated for MN, untreated AML | Combination venetoclax and azacitidine | CR flow MRD negative | Yes | CEBPA, STAG2, TET2, TP53, WT1 | 46,XY[20] | n/a | n/a |
| AML-DBP-18 (2) (biological duplicate) | 76 | Acute myeloid leukaemia, myelodysplasia-related | Refractory | gilteritinib | no response | Yes | CTCF, DNMT3A, RUNX1, SF3B1, TET2, TET2, TET2, FLT3-ITD | 46,XY[20] | n/a | n/a |
| AML-DBP-193 | 78 | Acute myeloid leukaemia, myelodysplasia-related, post-MPN | Refractory | Combination venetoclax and decitabine | no response | Yes | ASXL1, BCOR, CALR, NRAS, NRAS, and RUNX1 | n/a | n/a | n/a |
| AML-DBP-197 | 71 | Acute myeloid leukaemia with other defined genetic alterations | Untreated | Combination venetoclax and azacitidine | non-evaluable for response | Yes | NRAS, TP53, TP53 | 45,XX,-3,del(5)(q13q33),-7,+8,del(12)(p13p11.2)[20] | 93 | positive |
| AML-DBP-200 | 74 | AML with defining genetic abnormalities | Untreated | 7+3+midostaurin | CR flow MRD negative | Yes | NPM1 and FLT3-ITD | 46,XY[16] | 87 | positive |
| AML-DBP-202 | 67 | Acute myeloid leukaemia, myelodysplasia-related | Untreated | Combination venetoclax and azacitidine | CR flow MRD positive | Yes | NF1 | 44,XX,-3,der(5;17)(p10;q10),add(9)(p24),-15,add(19)(p13.3),-21,+r,+mar[20] | 28 | partial |
| AML-DBP-204 | 72 | Acute myeloid leukaemia, myelodysplasia-related | Untreated | Combination venetoclax and azacitidine | CRi flow MRD positive | Yes | ASXL1, IZKF1, NRAS, PTPN11, RUNX1 | 46,XX[20] | 11 | negative |
| AML-DBP-199c | 73 | Acute myeloid leukaemia, myelodysplasia-related | Previously treated for MN, untreated AML | no treatment received (supportive care only) | non-evaluable for response | No | n/a | 44,XY,add(6)(q2?2),-7,-18,del(19)(p13.1),del(20)(q11,2q13.3[cp3] | 46 | positive |
| AML-DBP-205 | 87 | Acute myeloid leukaemia, myelodysplasia-related | Untreated | gilteritinib | non-evaluable for response | Yes | FLT3-ITD, KMT2A-PTD, DNMT3A, RUNX1, TET2, WT1 | 46,XY[20] | 100 | positive |
| AML-DBP-206 | 76 | AML with defining genetic abnormalities, prior antecedant MPN | Untreated | dec/ven/navitoclax on clinical trial | CR | Yes | GNB1, JAK2, PTPN11, WT1 | 45,XX,inv(3)(q21q26.2),-7[16]/46,XX[4] | 69 | subset |
| AML-DBP-207 | 76 | Acute myeloid leukaemia, myelodysplasia-related | Untreated | Combination venetoclax and decitabine | non-evaluable for response | Yes | ASXL1, ETV6, EZH2, KRAS, NRAS, NRAS, RUNX1, TET2, TET2 and WT1 | 46,XY,-7,+8[18]/47,idem,del(9)(q13q22),+21[2] | 95 | dim |
| AML-DBP-209 | 73 | Acute myeloid leukaemia, myelodysplasia-related | Untreated | Combination venetoclax and decitabine | non-evaluable for response | Yes | DNMT3A, GNAS, TP53, TP53 | 46,XX,del(5)(q22q33)[2]/49-51,idem,add(11)(q21),+22,+3-4r[cp6]/52-54,idem, | 60 | variable |

| | | | | | | | | | | |
|---|---|---|---|---|---|---|---|---|---|---|
| AML-DBP-212 | 85 | Acute myeloid leukaemia, myelodysplasia-related | Untreated | Combination venetoclax and azacitidine | CR flow MRD negative | Yes | ASXL1, ATM, JAK2, KRAS, RIT1, and TET2 | 46,XY,?+Y,add(3)(p25),-4[cp2] | 71 | dim |
| AML-DBP-213 | 51 | Acute myeloid leukaemia with other defined genetic alterations | Untreated | CPX-351 | CR flow MRD negative | Yes | MYC expression | 47,XY,+r[12]/48,XY,+2r[8].ish r(MYC amp)x1~2[5] | 31 | dim, partial |
| AML-DBP-215 | 63 | Acute myeloid leukaemia with other defined genetic alterations | Relapsed/ Refractory | IS-free haplo BMT on clinical trial | no response | Yes | CEBPA, CEBPA, ETV6, ETV6, GNAS | 46,XX,t(12;15)(p11.2;q26)[12]/46,XX[cp8] | 72 | partial |
| AML-DBP-218 | 61 | AML with defining genetic abnormalities, with prior cytotoxic chemotherapy | Untreated | 7+3 | CR flow MRD negative | Yes | FLT3 I836del, FLT3 D835E, NRAS | -20,+21,+22[11]/62~69,idem,+X,+11,del(11)(p11.2),+1~2mar[7]/46,XX[2]. | 74 | variable |
| AML-DBP-220 | 79 | Acute myeloid leukaemia, myelodysplasia-related | Refractory | BXCL701 on clinical trial | no response | Yes | NRAS, RUNX1, RUNX1, RUNX1, SRSF2, TET2 | 46,XY | 33 | dim |
| AML-DBP-223 (1) | 62 | Acute myeloid leukaemia with defining genetic abnormality | Untreated | 7+3+midostaurin | CR flow MRD negative | Yes | FLT3-ITD, NPM1, TET2, TET2 | 46,XY[20] | 97 | positive |
| AML-DBP-224 | 70 | MDS/MPN | Relapsed | no treatment received (supportive care only) | no response | Yes | ASXL1, MPL, SRSF2 and TET2 | 46,XY[20] | 24 | variable |
| AML-DBP-223 (2) (biological duplicate) | 62 | Acute myeloid leukaemia with defining genetic abnormality | Untreated | 7+3+midostaurin | CR flow MRD negative | Yes | FLT3-ITD, NPM1, TET2, TET2 | 46,XY[20] | 97 | positive |
| AML-DBP-229 | 63 | AML with monocytic differentiation | Untreated | 7+3+midostaurin | CR flow MRD negative | Yes | ASXL1, FLT3 D835H and U2AF1; p210 BCR-ABL 0.0646%-0.8379% (extremely small subclone) | 46,XY,del(9)(q13q33)[13]/45,X,-Y,t(9;22)(q34;q11.2)[5]/46,XY[2].nuc ish(ABL1,BCR)x2[100] | 92 | positive |
| AML-DBP-233 | 53 | ACUTE MYELOID LEUKEMIA WITH INV(16)(p13.1q22) (ICC). ACUTE MYELOID LEUKEMIA WITH CBFB::MYH11 FUSION (WHO 5th ed). | Untreated | 7+3 + Gemtuzumab | CR flow MRD negative | Yes | KIT, NF1 and NRAS | 46,XX,inv(16)(p13.1q22)[20].nuc ish(PML,RARA)x2[100] | 51 | variable |
| AML-DBP-236 | 41 | ACUTE MYELOID LEUKEMIA WITH NPM1 MUTATION (WHO5, ICC). | Untreated | 7+3+midostaurin | CR flow MRD negative | Yes | DNMT3A, FLT3 D835Y, KRAS, NPM1 | 46,XY[cp20] | 55 | positive |
| AML-DBP-237 | 58 | ACUTE MYELOID LEUKEMIA WITH NPM1 MUTATION (WHO 5th ed.) ACUTE MYELOID LEUKEMIA WITH MUTATED NPM1 (ICC) | Untreated | 7+3+venetoclax on clinical trial | CR flow MRD negative | Yes | DNMT3A, IDH1, NPM1 and FLT3-ITDx2 (<0.01 VAF) | 46,XX[6] | 56 | dim |
